# Transcriptomic profiling of the embryonic *C. elegans* intestine with single-cell resolution

**DOI:** 10.64898/2026.05.20.726538

**Authors:** Jessica L Hill, Justin P Ellis, Robert TP Williams, Anthony Apodaca, Ambika Basu, Andrew Moore, Erin Osborne Nishimura

## Abstract

At a mere 20 cells, the *Caenorhabditis elegans* intestine regulates metabolism, energy homeostasis, host defense, yolk production, and genetic aging, all while dynamically responding to its environment. How the intestine develops to carry out these disparate functions is unknown, and how cells differ along the length of the intestine is unclear. To address these questions, we performed single-cell RNA sequencing (scRNA-seq) on FACS-enriched intestinal cells from mixed-stage *C. elegans* embryos. The resulting single-cell transcriptomes of 974 cells organized into 13 clusters, suggesting a diversity of cell types and states. We used two post hoc approaches to ascribe identities to each cluster. First, genes with known developmental timing in early-, mid-, and late-stages were used to place clusters in time, and smiFISH microscopy was used to fine-tune the assignments. Second, the eight late-stage clusters were assessed for their region of origin. To assign these clusters to anatomical regions, we identified marker genes for each cluster and assessed their expression along the anterior-to-posterior length of the intestine using smiFISH microscopy. Genes associated with growth and cell division were expressed in early stages, whereas genes associated with immune responses and metabolism were expressed later. Genes associated with biotic responses and RNA metabolism were the most likely to vary across the intestine’s anterior-posterior axis. Finally, perturbation of anterior-localized intestinal transcripts more robustly affected intestinal function compared to central or posterior-localized genes. Overall, this research illustrates the intrinsic heterogeneity across the 20 cells of the embryonic intestine and sets the stage for future works aimed at understanding cell-specific intestinal responses to diet and the environment.

**ARTICLE SUMMARY:** We investigate how the *Caenorhabditis elegans* intestine develops specialized functions on a spatiotemporal scale. We used single-cell RNA-sequencing to analyze embryonic intestinal cells and identify 13 distinct clusters. Combining gene expression analysis with microscopy, we assigned clusters to developmental stages and anatomical regions. Clusters associated with early intestine development express genes linked to growth and cell division, while later-stage clusters express genes involved in metabolism and immune responses. Genes varied across the intestine’s anterior-to-posterior axis, and disrupting anterior-specific genes produced stronger functional effects. These findings reveal previously unrecognized intestinal diversity and provide insight into how intestinal cells specialize during development.

## INTRODUCTION

Organogenesis is a complex process during which differentiating cells and germ layers develop into organs through concerted communication between gene networks and signaling pathways (Chevalier 2022). Our understanding of organogenesis is aided by the study of model organisms, especially those with simplified body plans and defined gene regulatory networks (Mango 2009; Irion and Nüsslein-Volhard 2022). The *Caenorhabditis elegans* intestine serves as a powerful model of organogenesis. At only 20 cells, the intestine arises clonally from a single E (endodermal) cell, is specified by a well-studied network of interconnected GATA transcription factors, and takes 24 hours to form (McGhee 2007; McGhee 2013; Maduro 2017; Dimov and Maduro 2019). Yet, this organ performs a wide diversity of functions: digestion, insulin regulation, immune response, detoxification, glycogen storage, yolk production, and genetically encoded aging. A major question is whether these functions are spatially organized across the length or breadth of the intestine and how such spatial differences might arise during organogenesis.

A greater understanding of the spatial organization of the *C. elegans* intestine would require high-resolution analysis. Fortunately, defining *C. elegans* intestinal development at the single-cell level is an achievable goal aided both by the invariant nature of *C. elegans* development, in which cell lineages have been fully mapped from fertilization to hatching (and beyond), and by its small size (Sulston and Horvitz 1977; Sulston et al. 1983). Intestinal cells arise over four mitotic events spanning the 8-cell to 100-cell embryonic stages. The resulting 20 cells then organize into either 2-cell or a 4-cell ring structures that link to form a tube connecting the pharynx at its anterior to the hindgut at the posterior (Dimov and Maduro 2019; Guan et al. 2025). This forms the gut or intestine of the worm. To date, our understanding of this developmental process has greatly benefited from research approaches at two different levels. The behaviors of individual genes and proteins have been studied at single-cell resolution using microscopy and genetic approaches (Hawkins and McGhee 1995; Fukushige et al. 1999; Bao et al. 2008; Goszczynski et al. 2016), while genomics assays have yielded insight into expression of the transcriptome, as a whole, from bulk embryos, worms, and intestinal tissues (Araya et al. 2014; Wiesenfahrt et al. 2015; Dineen et al. 2018; Kudron et al. 2018; Williams et al. 2023). Research in this manuscript aims to bridge these two capacities by assessing *C. elegans* intestinal gene expression at both genome-wide and single-cell levels.

Single-cell-resolution investigations of intestinal gene expression have been successful. These ventures include single-cell RNA-seq (scRNA-seq) assays performed on whole worms using Cel-seq2 (Cole et al. 2023), cellular dilution (Cao et al. 2017), or the commercially available 10x Genomics approach (Packer et al. 2019; Large et al. 2025). These studies, though aimed at assessing differences across organs, have revealed that transcriptomes of individual intestinal cells do vary. However, because these studies were aimed at assessing transcriptome differences across the whole organism, they did not enrich for intestinal cells specifically and obtained fewer intestinal cells than is ideal for clustering among the intestinal cells. It is important to keep in mind that although the intestine, at hatching, makes up 30% of the worm’s body mass, it is only 2% of its total cells, and this low number accounts for its low frequency in scRNA-seq assays.

In this study, we focused specifically on the intestinal lineage to characterize gene expression in developing intestinal cells at single-cell resolution. To do so, we first isolated populations of intestinal cells from dissociated mixed-stage embryos using FACS enrichment on an intestine-specific GFP marker strain. This was followed by 10x Genomics scRNA-seq on those intestinal cells to obtain single-cell, whole transcriptome expression profiles. The goal of this effort was to address two major questions. First, we asked how gene expression patterns of individual, developing intestinal cells changed throughout embryogenesis as they divided from the single E cell to all 20 later-stage cells. Second, we asked how the 20 cells of the late-stage intestine varied among one another. Such differences will yielded insights into how intestinal functions are spatially organized and how the process of organogenesis prepares the intestine for its myriad of roles, even before the worm hatches and begins to eat.

## MATERIALS AND METHODS

### *C. elegans* strains and maintenance

The JM149, MR142, and ERT60 worm strains were used for this study (**Table 1**). The wildtype strain N2 (Bristol) was used as a control. Worms were maintained as described (Stiernagle 2006) and were cultured at 20°C on NGM (3g NaCl, 17g agar, 2.5g peptone, 1mL 1M CaCl2, 1mL 5mg/mL cholesterol, 1mL 1M MgSO4, 25mL KPO4 buffer (pH 6) [108.3g KH2PO4 and 35.6g K2HPO4 in 1L H20], 1mL 10mg/mL Nystatin) plates seeded with the *Escherichia coli* (*E. coli*) strain OP50. Worm strains (JM149, MR142, ERT60, and N2) and *E. coli* OP50 were provided by the Caenorhabditis Genetics Center (CGC), which is funded by NIH Office of Research Infrastructure Programs (P40 OD010440). All strains are available upon request from the CGC.

**Table 1.** Strains used in this study are listed by name and genotype.

| Strain Name | Genotype |
| --- | --- |
| JM149 | *cals71[elt-2p::GFP::HIS-2B::unc-54 3'UTR + rol-6(su1006)]* |
| MR142 | *cdc-25.1(rr31) I; rrls1[elt-2::GFP + unc-119(+)]* |
| ERT60 | *jyls13[act-5p::GFP::ACT-5 + rol-6(su1006)] II* |
| N2 |  |

### *C. elegans* culture and preparation of dissociated embryonic *C. elegans* cells

To obtain mixed stage *C. elegans* embryos for dissociation and Fluorescence Activated Cell Sorting (FACS), large batches of gravid worms were reared as detailed in dx.doi.org/10.17504/protocols.io.8epv59zjng1b/v2. Briefly, *C. elegans* were grown on NGM plates seeded with *E. coli* OP50 at 20°C. Worms were synchronized twice to ensure gravid populations. Synchronization was performed by sacrificing worms with bleach solution to release mixed stage embryos, performing multiple washes to remove the bleach, incubating embryos in M9 buffer to hatch worms under starved conditions and synchronize their growth and arrest in the L1 stage (Stiernagle 2006).

100,000 2x synchronized embryos were seeded onto 20 total 150 mm NGM plates at a density of about 5,000 embryos per plate. Worms were cultured for 72 hr until gravid and harvested for mixed-stage embryos through bleach treatment. Embryos were then dissociated through enzymatic treatment with Chitinase and Pronase E, followed by mechanical disruption with a 21-gauge syringe needle. Post-dissociation, samples were kept on ice to improve viability. Sample concentrations were then determined by counting cells with the TC20 automated cell counter (Bio-Rad; 1450102). Samples were diluted as needed to a final sample concentration of 1×10^6^ cells/mL prior to running through the cell sorter. A detailed protocol for embryo dissociation is available on the protocols.io platform: https://dx.doi.org/10.17504/protocols.io.dm6gpbw9plzp/v1.

### FACS isolation of intestinal cells

Disassociated intestinal cells of the JM149 strain were enriched by FACS based off their GFP (Green fluorescent protein) signal that was intestine-specific due to *elt-2* promoter-driven GFP fluorescence. Propidium Iodide (ThermoFisher scientific; R37169) was used, per the manufacturer’s instructions, as a viability dye to separate live cells (PI^−^) from dead cells (PI^+^) prior to intestinal cell enrichment. As a sequencing control, dissociated JM149 embryonic cells were also run through the FACS analysis but were not gated at a specific GFP fluorescence level for collection (all cells control). Four biological replicates were captured for each group (intestinal cells and all cells control) for a total of eight samples (**Supplemental Figure 1**).

The BD FACS Aria III Cell Sorter was used with FACSDiva software version 6.1.3 to perform FACS enrichment of intestinal cells versus all cells. The cell sorter was cooled to 4°C prior to sample acquisition to improve cell viability. The following gating strategy was used to acquire intestinal cells: FSC-A vs SSC-A (intact cells), FSC-A vs FSC-H (single cells), PI-A vs SSC-A (live cells, PI^−^), FITC-A vs SSC-A (intestine cells, GFP^+^). Whereas to acquire the all cells control sample, the following gaiting strategy was used: FSC-A vs SSC-A (intact cells), FSC-A vs FSC-H (single cells), PI-A vs SSC-A (live cells, PI^−^) (**Supplemental Figure 1**). Samples were collected into a non-treated V-bottom plate (Greiner bio-one; 651182) for post-sort concentration. A total of 60,000 cells were captured for all samples. Samples were then concentrated by centrifugation (500 x g, 5 min, 4°C) to a final concentration of 1,200 cells/*μ*L for each sample. Samples were kept on ice until prepared for loading into the 10x Genomics Chromium Controller.

All cell sorting was done within the Colorado State University Flow Cytometry, Cell Sorting, and Single Cell Analysis Core Facility (RRID:SCR_022000) with assistance and guidance from core facility members.

### Sample preparation and sequencing

To perform single-cell RNA-seq, the 10x Genomics platform was chosen to perform the cell partitioning and library preparation. A total of eight samples (4 biological replicates for ‘intestinal cells’ and the ‘all cells’ control group) were sequenced, and these were processed in two separate batches (<u>batch 1</u>: intestinal cells rep 1 and rep 2 plus the all cells control rep 1 and rep 2; <u>batch 2</u>: intestinal cells rep 3 and rep 4 plus the all cells control rep 3 and rep 4).

Post-sort sample preparation was carried out for batch 1 samples by the CSU Genomics Core facility personnel and using their equipment (Chromium controller). For batch 2 samples, library prep and quality control were completed in lab (Chromium X). Prepared libraries were then sequenced at the at the CSU genomics core (batch 1) and CU Anschutz Medical Campus Gates Institute Genomics Core (batch 2).

We utilized the whole transcriptome Chromium Next GEM Single Cell 3’ Kit v3.1 (10x Genomics; PN-10000269) in combination with the Chromium Next GEM Chip G Single Cell Kit (10x Genomics; PN-10000127) and Dual Index Kit TT Set A (for Gene Expression Libraries) (10x Genomics; PN-10000215). All kits were used per the manufacturer’s instructions.

For GEM generation, samples at a concentration of 1,200 cells/*μ*L, were loaded into the Chromium Controller or Chromium X with a target capture between 6,000-9,000 cells. This concentration was chosen to maximize cells captured with reasonably low rates of multiplets. Following capture and lysis, cDNA was synthesized and then amplified. Amplified cDNA was then used to construct Illumina sequencing libraries. All sequencing libraries were pooled (for batch 1 and batch 2 respectively) to acceptable concentrations of around 10nM. Prepared libraries were sequenced on an Illumina NextSeq (batch 1) and NovaSEQ X (batch 2) platform to a depth of about 100 million reads per library.

### Data processing and analysis

Raw sequencing data was processed using the 10x Genomics Cell Ranger pipeline (v 8.0.1). For batch 1 samples, first Illumina base call (BCL) files were converted into FASTQ files. For batch 2 samples, the sequencing core provided the sample FASTQ files directly. Sequences were aligned to a custom-built transcriptome for *C. elegans* using Cell Rangers implementation of the STAR aligner (v 2.7.2a). The custom reference transcriptome was generated in Cell Ranger using available FASTA and GTF files (WBcel235) from Ensembl database and is included in the NCBI GEO Submission. Next, transcript abundance was quantified in each cell using Chromium barcodes. Finally, biological replicates for the ‘intestinal cell’ group and the ‘all cells’ control group were aggregated respectively into single datasets for ‘intestinal cells’ and ‘all cells’.

Analysis of processed scRNA-seq data was performed in R (v.4.5.1) using Seurat (v.5.4.0). 5,265 intestinal cells (GFP+) and 22,819 all cells (GFP+ and GFP-) passed Cell Range quality control filtering. Additional quality controls were performed in Seurat. Specifically, we filtered out cells that had either <200 or >5,000 unique genes expressed, >50,000 unique molecular index’s (UMIs), and/or >50% of reads mapping to the mitochondria. This resulted in a total of 2571 <u>intestinal cells</u> and 18,738 <u>all cells</u>. From the final intestinal cell dataset, we quantified gene expression across 15,116 genes. From the final all cells dataset, this was across 18,590 genes.

To investigate expression patterns and cluster transcriptionally similar cells, we first identified highly variable genes and reduced dimensionality using principal component analysis (PCA). For the intestinal cell dataset, we selected 5 principal components (PCs) as the relevant dimensions. This was done by both assessing the plotted standard deviations of the principal components (elbow plot) and by using the Seurat-implemented permutation-based test (JackStraw analysis and plot). We used PC loadings as input for UMAP (uniform manifold approximation and projection) embedding to reduce the dimensionality of the data to two for visualization purposes. Clustering resolution was determined using the R package clustree (v.0.5.1), which allows for visualization of cell cluster assignment across different resolutions. After clustering transcriptionally similar cells, we then identified and analyzed genes that were characteristic of each identified intestinal cluster. We used the non-parametric Wilcoxon Rank Sum test, as implemented in Seurat’s FindAllMarkers() function, to identify genes whose expression was enriched in specific clusters. This is effectively a two-sided test, where we first asked, prior to actually testing, that we restrict testing to only genes that 1) can be detected in at least 25% of cells from either group being compared (min.pct=0.25), that way we remove sparse/noisy genes, and 2) have a minimum fold difference of 1 between the either groups being compared (logfc.threshold=1). Of the genes that were actually tested and had a significant adjusted p-value (from multiple testing correction using Bonferroni method), we took the positive genes representing cluster enrichment (not depletion) into downstream gene ontology enrichment analysis.

We performed Gene Ontology enrichment analysis of cluster enriched genes (“cluster marker genes”) using WormBase’s (Davis et al. 2022) tissue enrichment analysis tool and clusterProfiler (v 4.18.4) (Yu et al. 2012; Wu et al. 2021; Xu et al. 2024). For clusterProfiler, gene symbols were converted to ENTREZ identifiers and analyzed using compareCluster() with enrichGO against either a background consisting of all cluster-associated genes or a background relative to gene sets derived from clusters belonging to remaining groups of interest (i.e., anterior clusters versus central and posterior clusters). Then ‘biological processes’ GO terms were filtered using Benjamini-Hochberg adjusted p-values (q<0.05) and semantically redundant terms were collapsed.

### Single molecule inexpensive fluorescence *in situ* hybridization (smiFISH)

smiFISH was used as a complementary approach to determine mRNA abundance and localization in *C. elegans* embryos for select transcripts. Transcripts were probed throughout embryonic development using smiFISH as previously described (Tsanov et al. 2016; Parker et al. 2021). To generate smiFISH probesets, 12-24x 18nt DNA oligonucleotide probes complementary to the transcript of interest were annealed *in vitro* to fluorescently labeled secondary probes to create the transcript-specific probesets (**Supplemental Table 1**). The accumulation of multiple fluorescent probes hybridizing to a single mRNA molecule produces a discrete, punctate digital signal *in situ*.

For each target transcript, 12 to 24 primary probes were designed using the R script Oligostan (https://bitbucket.org/muellerflorian/fish_quant/src/master/Oligostan/). All primary probes were designed with a FLAP X extension for detection by the fluorescently labeled secondary probe. The secondary FLAP X probe was designed and ordered from Stellaris LGC with dual 5’ and 3’ fluorophore labeling. We utilized both Cal Fluor 610 (LGC Biosearch Tech.; 5-/3-CR610-1) and Quasar 670 (LGC Biosearch Tech.; 5-/3-Q670-1) labeled FLAP X probes, allowing for multiplexing within the same sample.

Mixed stage embryos were harvested and prepared as described in the ‘Basic Protocol 3’ of Parker et al. 2021 (Parker et al. 2021). Briefly, JM149 worms were grown on large (150 mm) NGM plates seeded with *E. coli* OP50 until gravid, then synchronized by bleaching. Isolated embryos were then allowed to hatch overnight in M9 and arrest in the L1 stage. Synchronized L1 worms were then seeded back onto large NGM/OP50 plates and maintained at 20°C until gravid. Once gravid, mixed stage embryos were collected from worms by bleaching. Mixed stage embryos were then fixed and hybridized with smiFISH probesets overnight. Embryos were then washed, stained with DAPI, and prepped for fluorescence microscopy.

Embryos were imaged using a GE DeltaVision Elite inverted widefield fluorescent microscope equipped with an Olympus PLAN APO 60X, 1.42 NA objective, an Insight SSI 7-color Solid State Light Engine, and the standard DeltaVision DAPI, FITC, mCherry, Cy5 polychroic filter set. SoftWorx software version 7.2.2 (Cytiva) was used to operate and acquire images. Image acquisition settings were optimized for each target transcript. We acquire images through the entire Z-stack at the longest wavelength before moving to the next longest wavelength, to prevent photobleaching of the more labile fluorophores at the red end of the spectrum. We use 0.2 μm Z-spacing between images. DeltaVision (softWoRx) deconvolution software was applied for representative images. All images were processed in the ImageJ distribution Fiji (Schindelin et al. 2012).

### RNA interference (RNAi)

Select genes showing distinct temporal (early, mid, or late) or anatomical (anterior, central, or posterior) expression were chosen for further evaluation using RNAi to assess their impact on intestinal development. Genes chosen for RNAi targeting were: *ugt-14* (early development (dev.)), *cpr-1* (late dev., anterior intestine (int.)), *y32f6a.5* (late dev., anterior int.), *c14c6.5* (late dev., anterior int.), *clec-56* (late dev., central int.), *endu-2* (late dev., central int.), and *pbo-4* (late dev., posterior int.). We used *E. coli* feeding strains engineered to produce double-stranded RNA cognate to each of the selected target transcripts. These RNAi feeding strains were obtained from the Ahringer RNAi feeding library (Kamath and Ahringer 2003).

To visualize intestinal phenotypes, we performed RNAi depletion in the following strains: JM149, MR142, ERT60, and N2 (see **Table 1**). JM149 and MR142 express GFP driven from the *elt-2* promoter allowing for visualization of intestinal cells. MR142 contains supernumerary intestinal cells due to defects in CDC25.1 phosphatase activity during embryogenesis (Bao et al. 2008; Hebeisen et al. 2008; Hebeisen and Roy 2008).

To treat worms with RNAi and observe acute effects of target transcript knockdown on intestine physiology, freshly starved worms were chunked onto 150 mm NGM/OP50 plates to grow for 72 hr at 20°C until gravid. Synchronized embryos were isolated from this mixed stage population using bleach treatment. Synchronized L1 worms were grown on RNAi feeding strains for 48 - 72 hr at 20°C and then harvested. L4 and young adult worms were collected in 1 mL tubes and washed two times with M9. Worms were fixed by treating with room temperature methanol for 5 min followed by acetone for 5 min. To remove fixative, worms were washed two times with room temperature M9 buffer. These fixed worms were mounted on slides for fluorescence imaging as described above except using a LUCPLAN FL 40X, 0.60 NA objective. Fixation was necessary to remove autofluorescent gut granules that would have impaired our observation of desired intestinal fluorescent markers.

For the negative control experiment, we utilized a feeding strain containing the L4440 empty vector control. For the positive control experiment and confirmation of RNAi efficiency, we utilized a feeding strain targeting POP-1. RNAi experiments are considered successful when POP-1 RNAi resulted in 100% embryonic lethality. All RNAi plasmids were sequence verified before experimental use. One biological replicate was performed for each RNAi target among each worm strain.

## RESULTS

### Embryonic *C. elegans* intestinal cells discretely cluster

Specification and development of the *C. elegans* intestine initiates during embryogenesis with inductive cues and maternally loaded transcription factors (TFs): SKN-1, SPTF-3, POP-1, and PAL-1 (Dimov and Maduro 2019). This leads to the activation of successive pairs of GATA TFs MED-1/-2, END-1/-3, and culminates with ELT-2/-7. Starting with a singular E (endoderm) cell, the Ea (anterior) and Ep (posterior) cells arise (McGhee 2007). This is followed by their division into Eal (left) and Ear (right), and Epl and Epr respectively. Following this and two more rounds of division, the intestine contains 16 cells. At this stage, the primordial intestinal cells begin to polarize and form the lumen. A final round of division then occurs to complete the 20-cell intestine.

The *C. elegans* intestine is broadly categorized into intestinal rings (int rings). Each ring is denoted by its proximity to the anterior end of the worm, spanning from rings one to nine. The first int ring is made up of four intestinal cells, while the other eight rings are made up of only two intestinal cells. These somatic cells are never replaced and thus serve the worm throughout its life span. This allegorical characterization of the *C. elegans* intestine has provided a framework for understanding its organization and function, albeit with some limitations.

Our current understanding of how the *C. elegans* intestine organizes its functions stem from regional differences in structure (Maduro 2017). For example, the anterior intestine, specifically int 1 and 2, are observed to have shorter microvilli (Sulston and Horvitz 1977). As well as, from observations of distinctive patterning of specific transcripts, such as *ges-1* (Schroeder and McGhee 1998), *cpr-1* (Britton et al. 1998; Schroeder and McGhee 1998; Santo et al. 1999), *spp-9* (Madhu et al. 2020), and *pbo-4* and *itr-1* (Sulston and Horvitz 1977; Beg et al. 2008; Bender et al. 2013). Additionally, we know that the LIN-12/Notch pathway drives the twist of int rings 2, 3, and 4 (Sulston and Horvitz 1977; Sulston et al. 1983; Hermann et al. 2000; Neves and Priess 2005). Taken together, these previous data support a heterogenous model of intestine function, where intestine specific programs are carried out in a cell and/or region-specific manner. However, it remains to be understood how the *C. elegans* intestine organizes its transcriptome on an individual cell level. To begin to address this, we employ scRNA-seq of mixed stage embryonic intestinal cells using the 10x genomics platform, allowing exploration of the developing intestine (**Figure 1a**).

**Figure 1.**
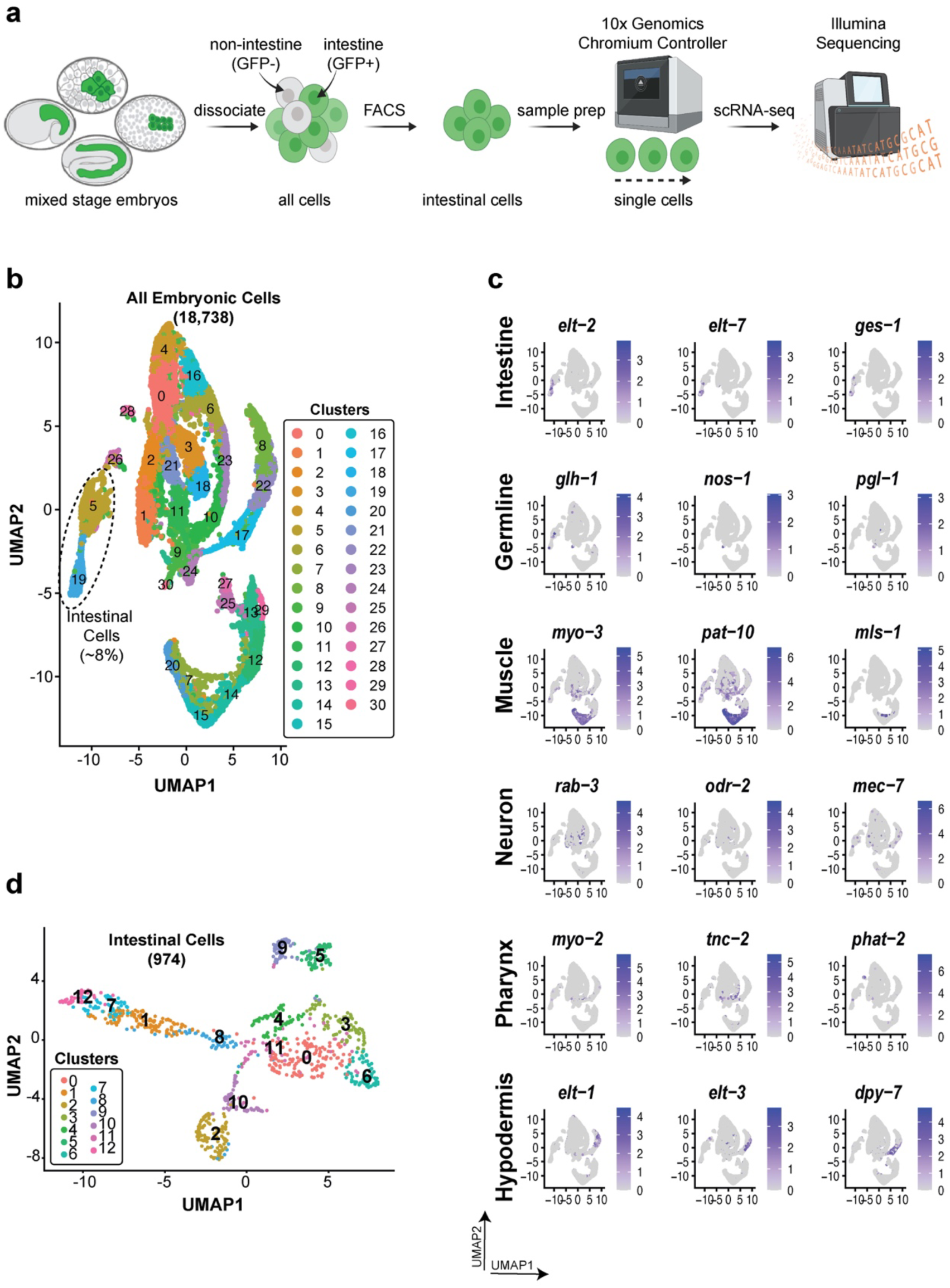
Characterization of the embryonic *C. elegans* intestinal cell transcriptome. **a)** Schematic representation of the approach to measure embryonic intestinal cell transcriptomes. **b)** UMAP embedding of scRNA-seq data from a mixed stage population of embryonic *C. elegans* cells. Intestinal cells cluster away from other tissue types and represent about 8% of total cells. **c)** Feature scatter plots displaying expression of major tissue marker genes from whole embryo dataset. **d)** UMAP embedding of scRNA-seq data from a mixed stage population of *C. elegans* intestinal cells. Thirteen distinct clusters are visualized.

Experimental datasets included the ‘intestinal cells’ group and the ‘all cells’ control group, each containing four biological replicates. 5,265 total ‘intestinal cells’ and 22,819 total cells from the ‘all cells’ control group passed Cell Range quality control filtering. Afterwards, additional quality control steps were implemented in Seurat. To remove potentially dead/dying cells or doublets, we filtered out cells having <200 or >5,000 unique genes expressed, >50,000 unique molecular index’s (UMIs), and/or >50% of reads mapping to the mitochondria. This resulted in a total of 2571 ‘intestinal cells’ and 18,738 ‘all cells’.

A final dataset of 18,738 cells for the all cells control group (**Figure 1b**) yielded 18,590 genes for quantification of differential expression. First, we identified highly variable genes and reduced dimensionality using PCA. Using the first 20 PCs, given their explanation of variability in this large dataset, cells were then clustered based on their transcriptional similarity. From the final dataset of 974 intestinal cells (**Figure 1d**), we quantified gene expression across 15,116 genes. For this dataset, we used the first 5 PCs, as they explained most of the variability in the data. As we had two batches of samples collected over time (rep1 and rep2, then rep3 and rep4) we wanted to look for the presence of batch effects in our data (**Supplemental Figure 2a-c and 3a-c**). There wasn’t a strong presence in the all cells control group dataset; however, there was a notable presence in the intestinal cell dataset. As such, we implemented the Anchor-based CCA integration method in Seurat to allow for correction of batch effects (**Supplemental Figure 2d-f and 3d-f**). Post clustering, we performed differential expression analysis to identify marker genes that distinctly characterize these integrated clusters (**Supplemental Table 2** (all cells) **and Supplemental Table 3** (intestine)); this will be discussed in the following section.

From the all cells control dataset, we were interested in determining how many tissue types were represented based on the expression of known marker genes. Broad classification of all the major tissue types was found (Intestine (*elt-2, elt-7, ges-1*), Germline (*glh-1, nos-1, pgl-1*), Muscle (*myo-3, pat-10, mls-1*), Neuron (*rab-3, odr-2, mec-7*), Pharynx (*myo-2, tnc-2, phat-2*), and Hypodermis (*elt-1, elt-3, dpy-7*)) (**Figure 1c**). Unsurprisingly, intestinal tissue represented roughly 8% of the total cells from this population, highlighting the necessity for enrichment of this particular cell type in order to generate a robust tissue specific dataset.

From the intestinal cells dataset, we found 13 unique clusters to describe the sequenced embryonic intestinal cells (**Figure 1d**). We recognize that this number of clusters does not directly correlate with the total number of intestinal cells (20) or intestinal rings (9). This suggests that more than just anatomical organization is driving how embryonic intestinal cells are clustering. Evaluated tissue marker genes for this intestine specific dataset are shown in **Supplemental Figure 4** and highlight a low level of contamination, as well as good representation of known intestinal tissue markers (*elt-2, elt-7, ges-1*) throughout the majority of the cells in this dataset. This is good practice to check, as animal dissociation and FACS enrichment of cells is not perfect. Additionally, it was important for us to try and remove disparate clusters, allowing for more subtle difference among intestinal cells to come through.

### Distinctive patterns of gene expression drive intestinal cell heterogeneity

We identified 13 distinct intestinal cell clusters in our dataset. To explore the driving forces behind why these cells cluster as they do, we visualized genes characteristic of each cluster. To do this, we first performed differential expression analysis and identified genes that are enriched in a given cluster (aka “cluster marker gene”). We utilized the Wilcoxon Rank Sum Test as implemented in Seurat’s ‘FindAllMarkers()’ function and pipeline. Effectively a two-sided test, we first asked, prior to testing, that we restrict testing to only genes that can be detected in at least 25% of cells from either group being compared, so as to remove sparse/noisy genes; second, we asked that we restrict testing to only genes that have a minimum fold difference of 1 between the either groups being compared. Of the genes actually tested and having a significant adjusted p-value from multiple testing correction using Bonferroni method, we took the positive genes representing cluster enrichment, not depletion. For clarification, although identified cluster marker genes are found to be significantly enriched in a given cluster, they may also be expressed to a lesser extent in other clusters.

Armed with this list of cluster marker genes, we made a heatmap of top marker genes for each of our identified clusters (**Figure 2**). Here, marker genes were filtered based on statistical significance (adjusted p-value<0.01) and then selected by ranking on average log2FC. For orientation, the heatmap displays genes as rows and cells in a given cluster as columns. Immediately, we can appreciate the distinctive patterning of marker genes across our intestinal clusters. Additionally, we can further appreciate how variable the cluster sizes are, corresponding to the number of cells in that given cluster (columns). This is also evident on the UMAP projection of our intestinal cells, but it becomes more stark in the heatmap. Importantly, there are clear chunks of enriched gene sets or groups that demarcate specific intestinal cell clusters.

**Figure 2.**
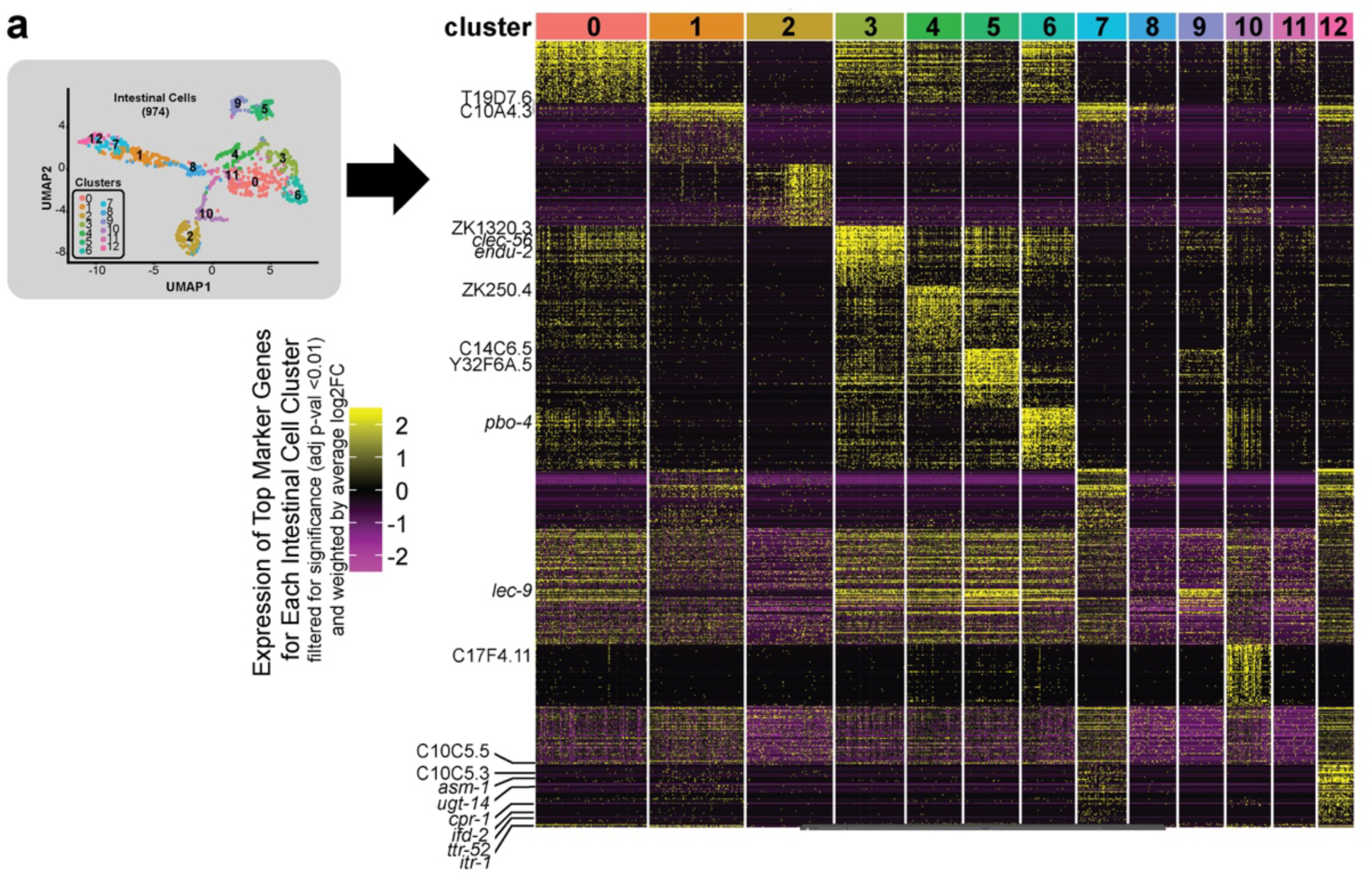
Di=erential gene expression across identified *C. elegans* intestinal clusters. **a)** (Left) UMAP embedding of scRNA-seq data from a mixed stage population of *C. elegans* intestinal cells, with 13 distinct clusters visualized. (Right) Expression heatmap of top marker genes for each intestinal cell cluster (gene = row x cluster/cells = column). A total of 64 genes are represented here. These marker genes were identified during diVerential expression analysis and filtered to retain significantly enriched genes (adjusted p-value<0.01), then for each cluster the top genes were selected based on highest average log2FC. Some of the identified cluster marker genes were then used for additional downstream analysis, of which they are highlighted on the left side of the heatmap.

While **Figure 2** highlights genes specifically enriched in individual clusters, emphasizing cluster identity, we also wanted to assess genes with the largest expression variability across all clusters, emphasizing global transcriptional heterogeneity and relationships between clusters (**Supplemental Figure 5**). What stands out are groups of seemingly related clusters (i.e., 0, 4, 6, 3, 5 and 12, 1, 7) in addition to clusters that appear to be stand-alone (i.e., 2, 10, 8).

Given this treasure trove of intestinal cluster associated genes, we can next inspect what intestinal cells these clusters relate to *in vivo* using <u>s</u>ingle <u>m</u>olecule inexpensive fluorescence *in <u>s</u>itu* <u>h</u>ybridization coupled with fluorescence microscopy. We can also investigate cluster associated functions using Gene Ontology.

### Spatiotemporal assignment of intestinal clusters

With the identification of 13 discrete intestinal cell clusters, we wanted to determine when and where these clusters present *in vivo*. To explore this, we utilized JM149 mixed stage embryos—same developmental stages and worm strain used to generate our scRNA-seq datasets—and probed for select cluster marker genes. To determine which cluster marker genes to actually probe for, we first turned to our list of all identified cluster marker genes and manually searched the top hits for known intestinal annotations, as well as predicted intestinal annotations. We used well-known intestinal genes to help inform our dataset and analysis (i.e., *cpr-1, pbo-4, lec-9*), as well as annotated previously uncharacterized genes. Additionally, we were interested in investigating genes along both the temporal axis (over developmental time) and spatial axis (anatomical regions of the intestine). We utilized reported expression values from WormBase to estimate when in embryogenesis these target transcripts may be turning on and off. Altogether, we identified a total of 19 targets to probe with smiFISH (**Table 2**). While all of these transcripts were surveyed, not all are represented in the main figures, as such their data can be found in **Supplemental Figures 6-8**.

**Table 2.**
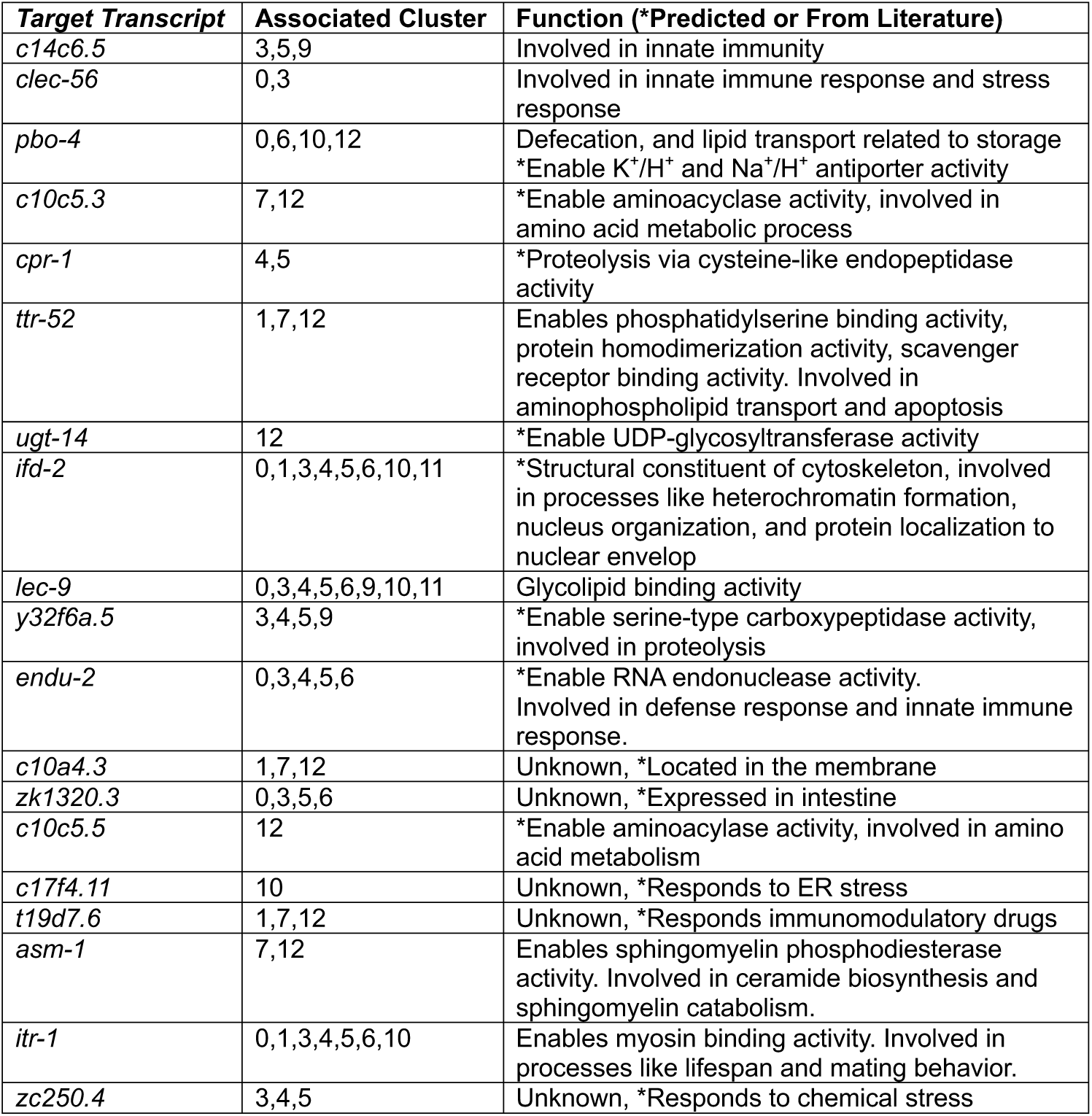
All intestinal cluster marker genes evaluated microscopically using smiFISH.

It is important to note that because we utilized the JM149 strain (see **Table 1**, *elt-2p*::GFP) for both scRNA-seq and smiFISH experiments, which allows for visualization of intestinal cell nuclei with GFP, we can only visualize intestinal cells as early as the *elt-2* promoter becomes activated and fluorescent protein accumulates (around the 64 cell stage (E^8^) (Wiesenfahrt et al. 2015; Maduro 2017)). Given that, we found that when we looked across embryogenesis, binning embryos into either early intestine development (64 cell stage, E^8^) (**Figure 3b**), mid intestine development (comma (E^16^), and 2-fold stage (E^20^)) (**Figure 3c**), or late intestine development (3-fold stage, E^20^) (**Figure 3d**), we could associate identified intestinal clusters with these developmental time points (**Figure 3**). Specifically, smiFISH target *ugt-14* (**Figure 3b**) was found to be expressed in the 64-cell stage but not any of the later stages. From our UMAP, *ugt-14* was associated with cluster 12, allowing us to associate cluster 12 with early intestine development. Other representative genes from cluster 12 were identified (*acp-2, end-1, c10c5.4*) and their expression patterns cross checked from reported studies on WormBase. For the smiFISH target *t19d7.6*, we found it was expressed in the comma and 2-fold stages only (mid intestine development), and was associated with clusters 1, 7, and 12 (**Figure 3c**). Other representative genes from these clusters (1, 7, 12) and cluster 4 were identified (*t21e8.4, c10a4.3* (**Supplemental Figure 7**)*, klo-1*) and cross checked, allowing us to associate these clusters with mid intestine development. Finally, the smiFISH target *cpr-1*, was observed in the 3-fold stage only. From our UMAP, *cpr-1* was associated with clusters 4 and 5. When we checked the timing of expression for other genes from these clusters, we found that other late intestine development genes (i.e., *lec-9* (**Supplemental Figure 7**)*, clec-56* (**Supplemental Figure 6**)*, c14c6.5* (**Supplemental Figure 6**)) were expressed more broadly. As such, we were able to associate clusters 0, 3, 4, 5, 6, 9, 10, and 11 with late intestine development. It is interesting to note the bit of overlap between these adjacent developmental windows and thus clusters, highlighting the presence of more modest differences in these clusters.

**Figure 3.**
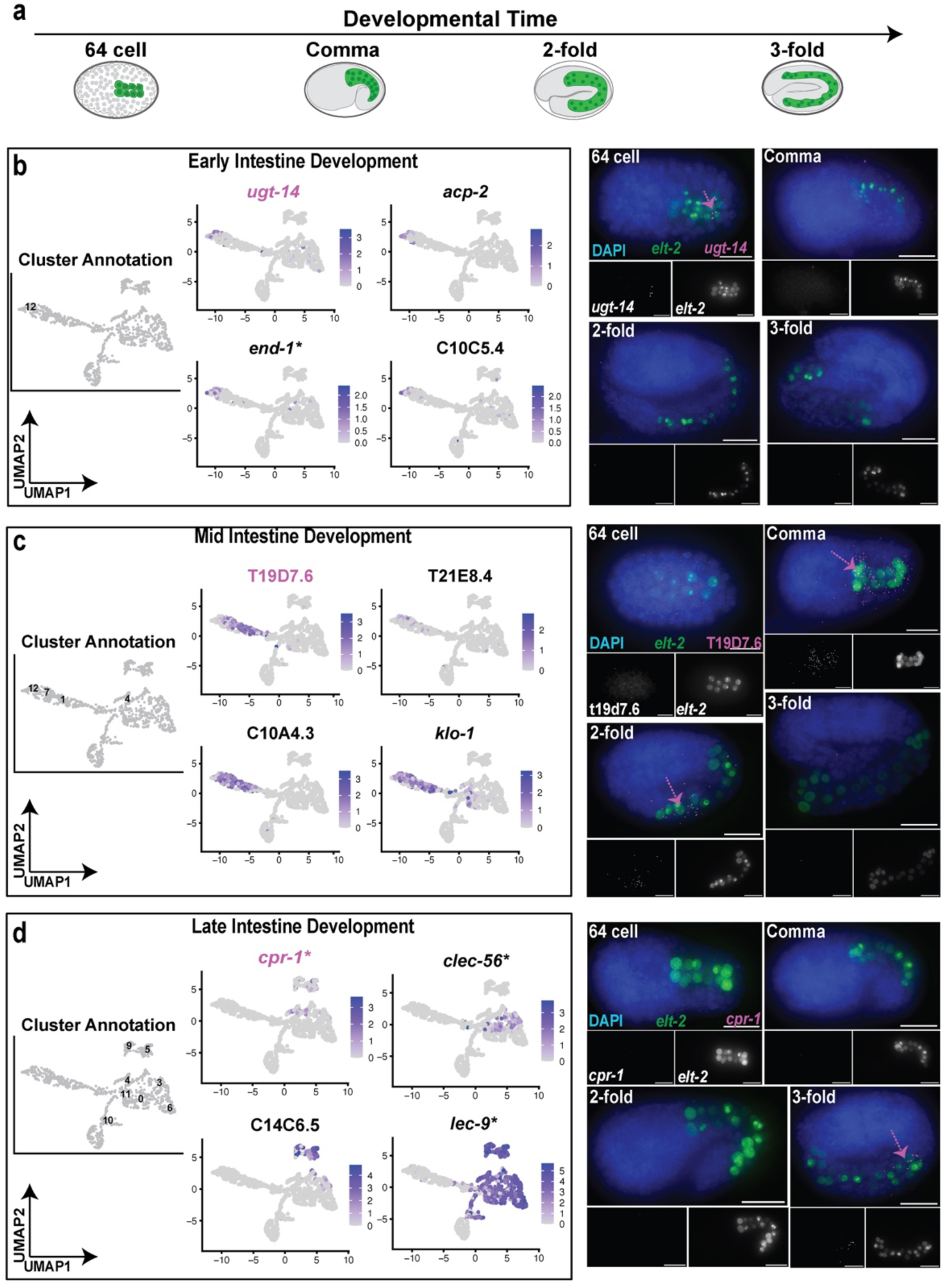
Distinct transcripts resolve identified embryonic *C. elegans* intestinal clusters based on developmental timing of gene expression. a) Schematic representation of sampled embryonic developmental time points using single molecule inexpensive Fluorescence *in situ* hybridization (smiFISH), with the intestine highlighted in green. **b)** Cluster marker genes associated with <u>early intestine development</u>. From left to right, UMAP embedding of intestinal cluster annotations for selected transcripts, accompanied by feature scatter plots displaying their expression level and distribution among cells. Representative smiFISH images of highlighted gene in magenta, across embryonic developmental time points. Similarly, cluster marker genes associated with <u>mid intestine development</u> (**c**) and <u>late intestine development</u> (**d**) are shown. Genes denoted with a * indicate already having intestinal annotation.

We next wanted to annotate our identified intestinal clusters along the spatial axis (anatomical regions of the intestine). For this, we binned the *C. elegans* intestine into anterior, central, and posterior regions (**Figure 4a**). Using our smiFISH targets *c14c6.5* and *cpr-1*, we found that they were exclusively expressed in the anterior region of the intestine and that this corresponded with clusters 3, 4, 5, and 9 on our UMAP (**Figure 4b**). As such, we assigned these clusters as associated with the anterior intestine. In contrast, using our smiFISH targets *clec-56* and *endu-2*, we found that they were expressed within the central region of the intestine (the largest apparent region), corresponding with clusters 0, 3, 4, 5, and 6 on the UMAP (**Figure 4c**). We thus associated these clusters with the central intestine. Finally, using the smiFISH target *pbo-4*, we demonstrate that it is exclusively expressed within the posterior region of the intestine, affiliated with clusters 0, 6, and 10; as such, we associated those clusters with the posterior intestine (**Figure 4d**). Of note, we were not yet successful in finding other target transcripts with exclusively posterior expression, hence pbo-4 is our only representative for that group. Also, we again highlight the apparent bit of overlap between proximal regions and related clusters, suggesting smaller differences in these clusters.

**Figure 4.**
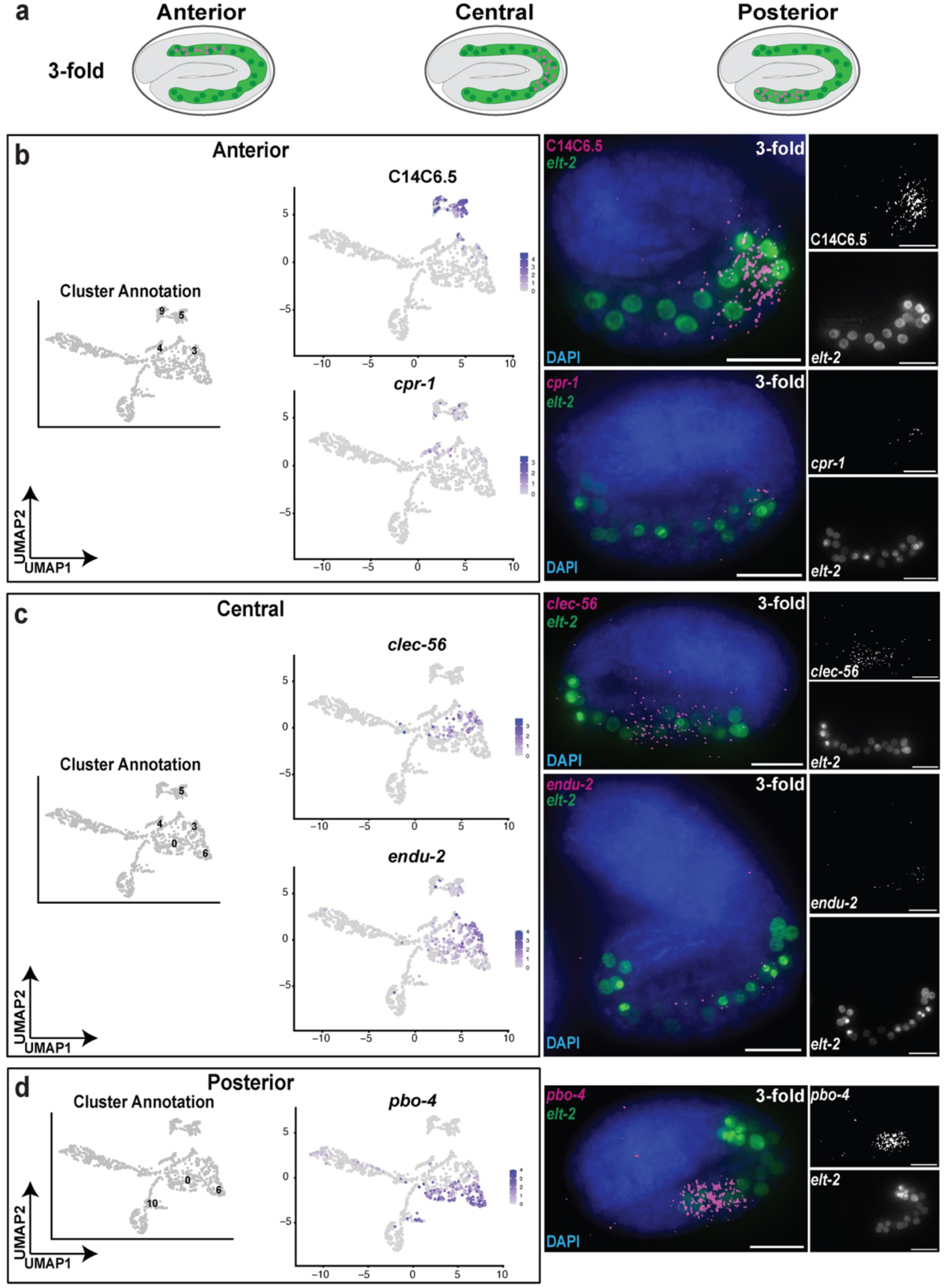
Distinct transcripts resolve identified embryonic *C. elegans* intestinal clusters based on anatomical region throughout the intestine. a) Schematic representation of sampled anatomical expression patterns (anterior, central, and posterior) using single molecule inexpensive Fluorescence *in situ* hybridization (smiFISH), with the intestine highlighted in green and the expression pattern highlighted in magenta. **b)** Cluster marker genes associated with the <u>anterior intestine</u>. From left to right, UMAP embedding of intestinal cluster annotations for selected transcripts, accompanied by feature scatter plots displaying their expression level and distribution among cells, followed by representative smiFISH images in the 3-fold stage. Similarly, cluster marker genes associated with <u>central intestine</u> (**c**) and <u>posterior intestine</u> (**d**) are shown. Genes denoted with a * indicate already having intestinal annotation.

Taken together, these findings highlight that the identified intestinal cell clusters from our UMAP appear to be clustering based on developmental time on the x-axis and based on anatomical region within the intestine on the y-axis. It is intuitive that regional differences in intestinal anatomy and function would evolve in the later stage intestine prior to embryo hatching. It is also important to remember that this dataset represents the intestinal transcriptome throughout embryogenesis and affords a glimpse into the developing intestine. How the intestine may change and further specify upon exposure to food and the environment is likely more drastic and should be explored.

### Functional annotation of intestinal clusters

Now that we have annotated our intestinal cell clusters with temporal and spatial identities, we can further characterize them based on enrichment of function associated terms using gene ontology enrichment analysis. Towards this end, we explored intestinal cluster-associated marker genes to identify biological processes (BP) enriched within specific intestinal populations (**Figure 5**). GO enrichment analysis across all intestinal clusters revealed substantial functional heterogeneity (**Figure 5b**). To investigate the spatial organization of intestinal functions, we grouped clusters according to inferred anatomical region (anterior, central, posterior) and analyzed GO enrichment relative to the remaining anatomical regions (**Figure 5c**). This revealed region specific transcriptional programs. For example, anterior and central intestine regions appear enriched for immune defense response pathways, while the posterior intestine exhibits transitional and ribosome biogenesis related pathways (**Figure 5c**). Similarly, grouping intestinal clusters according to inferred developmental timing revealed distinct temporal programs throughout intestine development (**Figure 5d**). Specifically, early and mid-intestine development associated clusters appear enriched for biological processes related to chromosome organization, mitotic cell cycle progression, and cell division, consistent with proliferative intestinal precursor populations (**Figure 5d**). In contrast, later intestine development associated clusters show enrichment for ribonucleoprotein complex biogenesis, metabolic processes, and immune-associated pathways, suggesting progressive functional specialization during intestinal maturation. (**Figure 5d**).

**Figure 5.**
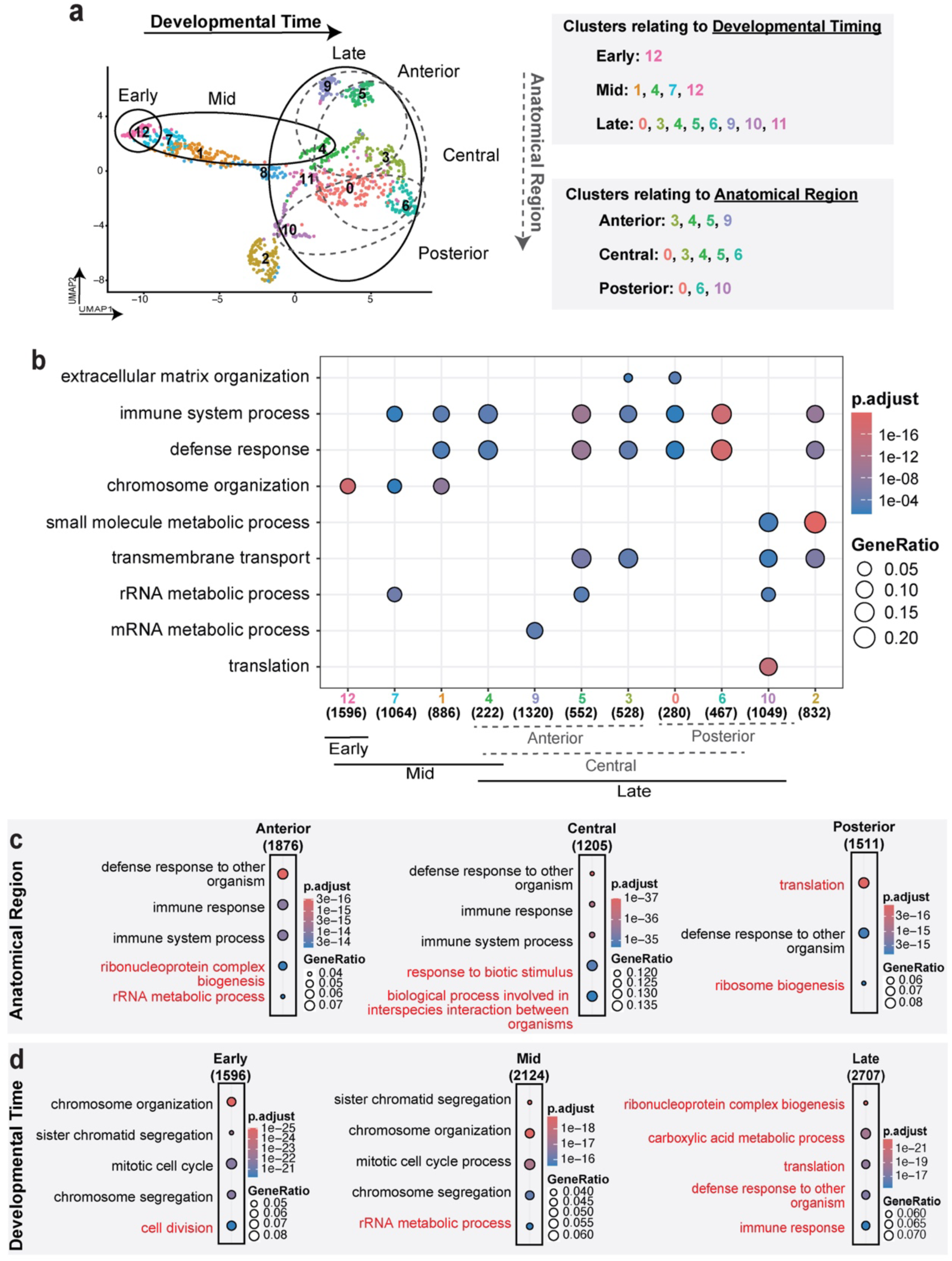
Integration of developmental, spatial, and functional identities across intestinal clusters. (**a**) UMAP projection of intestinal cell clusters, highlighting identified spatiotemporal organization with developmental timing on the x-axis and anatomical organization within the intestine along the y-axis (left). Cluster annotations are specified for clarification (right). (**b**) Gene ontology (GO) enrichment analysis of intestinal clusters, showing biological process terms. Gene symbols were converted to ENTREZ identifiers and analyzed using compareCluster() from clusterProfiler package (v 4.18.4) with enrichGO against a background consisting of all cluster-associated genes. GO terms were filtered using Benjamini-Hochberg adjusted p-values (q<0.05) and semantically redundant terms were collapsed. Dot size indicates gene ratio (number of genes from the cluster in the GO term/total number of genes in the cluster) and color represents adjusted p-value significance. Numbers in parentheses underneath cluster identities indicate the total number of genes in that cluster and thus used for analysis. (**c**) Representative enriched GO biological process terms associated with anatomically defined intestinal regions (anterior, central, posterior). Cluster-associated marker genes from clusters assigned to anterior, central, or posterior intestinal regions were grouped and analyzed using compareCluster() with enrichGO. Enrichement analysis was performed relative to background gene sets derived from clusters belonging to the remaining anatomical regions. Redundant terms were collapsed. Distinct regional transcriptional programs are highlighted in red. (**d**) Representative enriched GO biological process terms associated with developmental timing groups (early, mid, late). Cluster-associated marker genes from clusters assigned to early, mid, or late intestine development were grouped and analyzed as described above. Enrichement analysis was performed relative to background gene sets derived from clusters belonging to the remaining developmental stages. Redundant terms were collapsed. Distinct regional transcriptional programs are highlighted in red.

We were also interested in characterizing transcription factors associated with specific intestinal cell clusters. To do this, we utilized the curated wTF2.0 list of worm transcription factors (Reece-Hoyes et al. 2005) in addition to identified intestinal transcription factors pulled from the modEncode project (Nostrand and Kim 2013; Williams et al. 2023). Reasoning that these factors may contribute to establishment and maintenance of region-specific and temporally regulated intestinal identities. Unsurprisingly, we found a diversity of intestinal cluster associated transcription factors (**Supplemental Figure 9**). Most notably, we highlight that early intestine development associated clusters express developmental regulators (i.e., *end-1, odd-1*) and that there is a greater accumulation of stress-responsive (i.e., *pqm-1*), metabolic (i.e., *ets-4*), and tissue-patterning (i.e., *fkh-9*) transcription factors in the later intestine development group where we see anatomical segregation.

### Phenotypic evaluation of perturbation of spatiotemporally distinct intestinal transcripts

To investigate whether some intestinal transcripts (*ugt-14, cpr-1, y32f6a.5, c14c6.5, endu-2, clec-56, pbo-4*) identified as spatiotemporally distinct (see **Table 2**) contribute to intestinal development and function in any way, we performed RNA interference (RNAi) mediated knockdown in specific intestinal marker strains (JM149, ERT60, and MR142 (see **Table 1**)). These strains each report on a different aspect of intestinal physiology. The JM149 strain displays a wildstype intestine, with intestinal nuclei labeled with GFP driven under the *elt-2* promoter (transcriptional reporter). The MR142 strain displays a supernumerary intestine due to a gain of function mutation in the *cdc-25.1* gene, and has intestinal nuclei labeled with GFP driven under the *elt-2* promoter (transcriptional reporter). The ERT60 strain, displays a wild type intestine, with the intestinal lumen labeled with GFP driven by the ACT-5 protein (translation reporter), which gets incorporated in the intestinal villi.

Chosen RNAi targets span developmental time (early or late), but largely reflect different anatomical regions within the intestine. Using RNAi feeding libraries cognate to our target transcripts of interest, we fed synchronized L1 worms of each worm strain respective RNAi strains until they reached the L4 to young adult stage. We then characterized any embryonic lethal phenotypes present in laid embryos and collected worms for fixation and fluorescence microscopy to inspect changes in intestinal reporters and gross morphology.

In the *elt-2p*::GFP(JM149) worm strain (**Figure 6**), we found that egg laying capacity only appeared reduced by the knock down of *endu-2*, *ugt-14*, and *y32f6a.5*, with *endu-2* being the most drastic. These brood size phenotypes were also observed in the wild type N2 strain (data not shown). Interestingly, when we evaluated changes in intestinal fluorescence for this strain, we saw that knock down of anterior intestinal transcripts elicited the greatest changes in total intestinal fluorescence, followed by knock down of central intestine transcripts (**Figure 6a**). Likewise, when we evaluated changes in intestinal fluorescence for the ACT-5::GFP(ERT60) worm strain (**Supplemental Figure 10**) and the *elt-2p*::GFP(MR142) worm strain (**Supplemental Figure 11**), we found that perturbation of largely anterior intestine transcripts, and to a lesser extent central intestine transcripts, yielded the most drastic changes in total intestinal fluorescence. Notably, there were statistically significant increases in total intestinal fluorescence observed in the ACT-5::GFP(ERT60) strain when *cpr-1* was knocked down (**Supplemental Figure 10b**), and the *elt-2p*::GFP(MR142) strain when *y32f6a.5* was knocked down (**Supplemental Figure 11b**). Taken together, these data suggest that perturbation of the anterior intestine imparts a stronger impact on intestinal function than perturbation of other intestinal regions. This is understandable given that the anterior intestine may potentially be more important for setting up or establishing normal intestinal functions.

**Figure 6.**
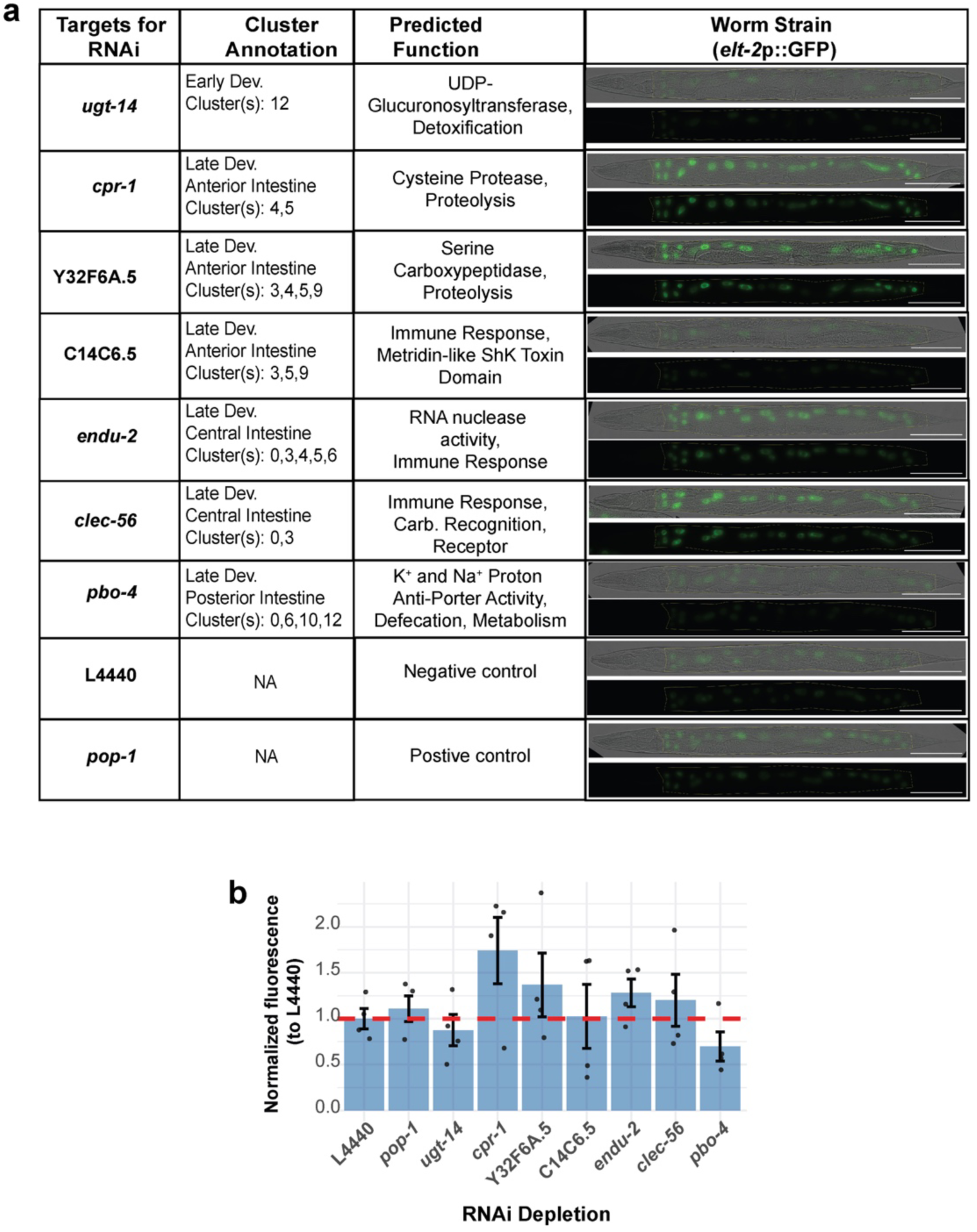
RNAi-mediated depletion of spatiotemporally identified transcripts in JM149 worm strain. (**a**) Representative brightfield and fluorescence images of L4 stage JM149 (*elt-2p*::GFP) worms after RNAi depletion of spatiotemporally distinct candidate smiFISH targets (*ugt-14*, *cpr-1*, *y32f6a.5*, *c14c6.5*, *endu-2*, *clec-56*, and *pbo-4*), in addition to treatment with the both negative (L4440) and positive (*pop-1*) control RNAi strains. (**b**) Quantification of intestinal *elt-2p*::GFP fluorescence after RNAi treatment. Fluorescence intensity was measured in individual animals, background corrected and then normalized to the mean fluorescence of the negative control (L4440). Bars represent mean normalized fluorescence ± SEM, and black dots indicate individual animals. The red dashed line marks the normalized L4440 negative control (1.0). Statistical significance was determined by comparing each RNAi treatment to the negative control using two-sided Wilcoxon rank-sum test (*p-value<0.05).

## DISCUSSION

In this investigation, we illustrate that the embryonic *C. elegans* intestine exhibits substantial transcriptional heterogeneity which organizes across both developmental time and anatomical space. Using FACS enrichment of intestinal cells coupled with scRNA-seq, we identified 13 discrete intestinal cell clusters from mixed-stage embryos, and using smiFISH, we assigned these clusters with developmental windows and anatomical regions. Together, these findings expand the current model of the embryonic intestine from a largely anatomical framework into one that includes dynamic and region-specific transcriptional specializations.

One important observation noted here is that intestinal cells did not cluster according to the canonical 20-cell or 9-ring organization of the mature intestine. Rather, their clustering reflected developmental state along one axis and anterior-posterior identity along the other axis. This suggests transcriptional identity within the intestine is not lineage fixed but rather emerges progressively during organogenesis. It also supports intestinal cells occupying translational molecular states rather than discrete identities. This model is consistent with the gradual establishment of intestinal polarity, lumen formation, and epithelial maturation observed during embryogenesis. These data also support the idea that regional intestinal specialization initializes prior to feeding. Here, we extend upon the prior knowledge of intestinal patterning with these transcriptome-wide observations. In particular, anterior and central intestine-associated clusters were found to be enriched for immune and defense-related pathways, whereas posterior intestine-associated clusters were seen to be enriched with ribosome biogenesis and translation processes, providing further evidence for a model in which distinct intestinal regions are molecularly primed for different functions prior to hatching.

By combining transcriptomic analysis with smiFISH across embryogenesis, a developmental progression of intestinal identities emerges. For instance, early development-associated clusters were found to express genes involved with chromosome organization, mitotic progression, and developmental specification. This is consistent with rapid proliferative events that generate the 20-cell intestine from the single E blastomere. Importantly, these early intestine clusters also expressed known developmental regulators like *end-1*, lending confidence to our temporal assignments. As embryogenesis progressed, mid and late development-associated intestinal clusters became enriched for metabolic and immune related pathways. One logical interpretation is that the intestine transitions from developmental programs focused on proliferation and morphogenesis towards those focused on physiological preparedness. Newly hatched larvae must interact with their environment and break down and absorb nutrients. As such it makes sense that these programs would accumulate over developmental time. Interestingly, and as noted previously, there was a noticeable amount of overlap between proximal clusters in our UMAP, suggesting that both temporal and regional transitions are continuous rather that sharply separated.

Another curious finding came from our RNAi experiments, which suggested that perturbation of anterior intestinal transcripts elicited stronger effects on intestinal physiology when compared to perturbations of central and posterior transcripts. Across multiple intestinal marker strains, knockdown of anterior genes via RNAi consistently produced the largest alterations in the intestinal reporters. Although many of these changes did not reach statistical significance, the consistency of the trend across strains is notable. One possible explanation is that the anterior intestine plays a more critical role in establishing or coordinating intestinal functions. Alternatively, anterior intestinal cells may simply experience greater functional demands. Nevertheless, many unknowns remain regarding how these regional identities are established and maintained. Further, it will be interesting to observe whether these embryonic regional identities persist into larval and adult stages and/or whether they are further remodeled following feeding and environmental exposure.

It is tempting to speculate on how the identified intestinal cluster-associated transcription factors may regulate underlying cluster-distinct or shared programs. For example, the observation of *end-1* and *odd-1* enrichment in early development associated clusters tracks with their known role as developmental regulators. In addition to this, we found enrichment of *pqm-1*, *ets-4*, and *fkh-9* (stress responsive and metabolic regulators) in later development associated clusters, which makes sense as this is where anatomical patterning presents. This cluster-specific transcription factors list suggests that distinct regulatory programs underlie the intestinal transitions that occur during embryogenesis. Further perturbation studies examining these factors in isolation or in combination will help tease apart how intestinal sub-functionalization is encoded transcriptionally and expand upon our understanding of intestinal GRNs.

Taken together, the work presented here establishes a framework for studying intestinal specialization during development, though several questions remain. One immediate direction will be to determine how transcriptional programs in the intestine change following hatching and exposure to diet, microbes, and environmental stressors. Because this study focused exclusively on *C. elegans* embryos, it is only capturing the earliest stages of intestinal sub-functionalization. Likewise, more extensive functional perturbation studies are necessary to determine whether regionally enriched transcripts contribute directly to programs like digestion, immunity, metabolism, etc. Ultimately, these findings support a model in which the embryonic intestine undergoes progressive transcriptional specializations prior to hatching, establishing spatially organized molecular programs that likely prepare the intestine for its diverse physiological roles throughout life.

## Accessibility Statement

The colors in microscopy images and plots were selected following guidelines for accessibility. We opted for magenta/green combinations in microscopy images.

## Data Availability Statement

Worm strains and plasmids used in this study are available upon request. Detailed protocols for embryonic dissociation and FACS isolation of intestinal cells are in the following protocols.io collection https://dx.doi.org/10.17504/protocols.io.5jyl895jdv2w/v1. Code for computational analysis and data visualization are available in the following GitHub repository https://github.com/jesshill/Celegans_Embryonic_Intestine_scRNAseq.git.

## Acknowledgments

We are thankful to everyone who assisted in small and large ways along the way and apologize if we forgot to specifically mention or cite colleagues here. We would like to say thank Dr. John Murray and his group for meaningful conversations on their experience with performing single cell RNA-seq in worms using the 10x genomics platform and subsequent data analysis. We thank everyone who has worked and currently works at the CSU Genomics Core Facility, making resources and expertise available to resident researchers. We thank the University of Colorado Anschutz Medical Campus Cancer Center Genomics Shared Resource Core Facility (RRID:SCR_021984). We thank Tai and Brooke Montgomery for housing and maintaining the *C. elegans* RNAi Collection (Ahringer) (RRID:SCR_017064), making these resources available. We thank WormBase ParaSite (Howe et al. 2017) and Ensembl Genomes (Kersey et al. 2014). We also thank WormBase and the Alliance of Genome Resources (Consortium et al. 2024). Finally, some strains were provided by the CGC, which is funded by NIH Office of Research Infrastructure Programs (P40 OD010440).

## Conflict of Interest or Competing Interests

The author(s) declare no conflict of interest. The authors have no competing or financial interests.

## Authors Contributions

Conceptualization: J.L.H., R.TP.W., E.O.N.; Methodology, formal analysis, and investigation: J.L.H., J.P.E., R.TP.W., A.A., A.B., A.M., E.O.N.; Writing, review, and editing: J.L.H., J.P.E., R.TP.W., A.A., A.B., A.M., E.O.N.; Supervision, project administration and funding acquisition: J.L.H., E.O.N.

