## Supplementary Material for "Transcriptomic profiling of the embryonic *C. elegans* intestine with single-cell resolution"

**Affiliations:**
[1]Department of Biochemistry and Molecular Biology, Colorado State University, Fort Collins, CO, 80523 USA

**ORCID IDs:** 0000-0002-3323-8379 (J.L.H.); 0009-0004-2531-3481 (J.P.E.); 0000-0002-3384-212X (R.TP.W.); 0009-0009-8599-0683 (A.A.); 0009-0005-5995-0371 (A.B.); 0000-0003-3267-669X (A.M.); 0000-0002-4313-4573 (E.O.N.).

**Competing interests:** The authors declare no competing or financial interests.

**Funding:** This work was supported by the National Institute of General Medical Sciences (R35GM124877), the National Institute of Diabetes and Digestive and Kidney Diseases (F32DK131636), the National Science Foundation (2143849), and the Boettcher Foundation (The Boettcher Webb-Waring Award).

**Running Title:** scRNA-seq of the *C. elegans* embryonic intestine

**Keywords:** intestine, *C. elegans*, embryogenesis, scRNA-seq, gut

**Figure S1**

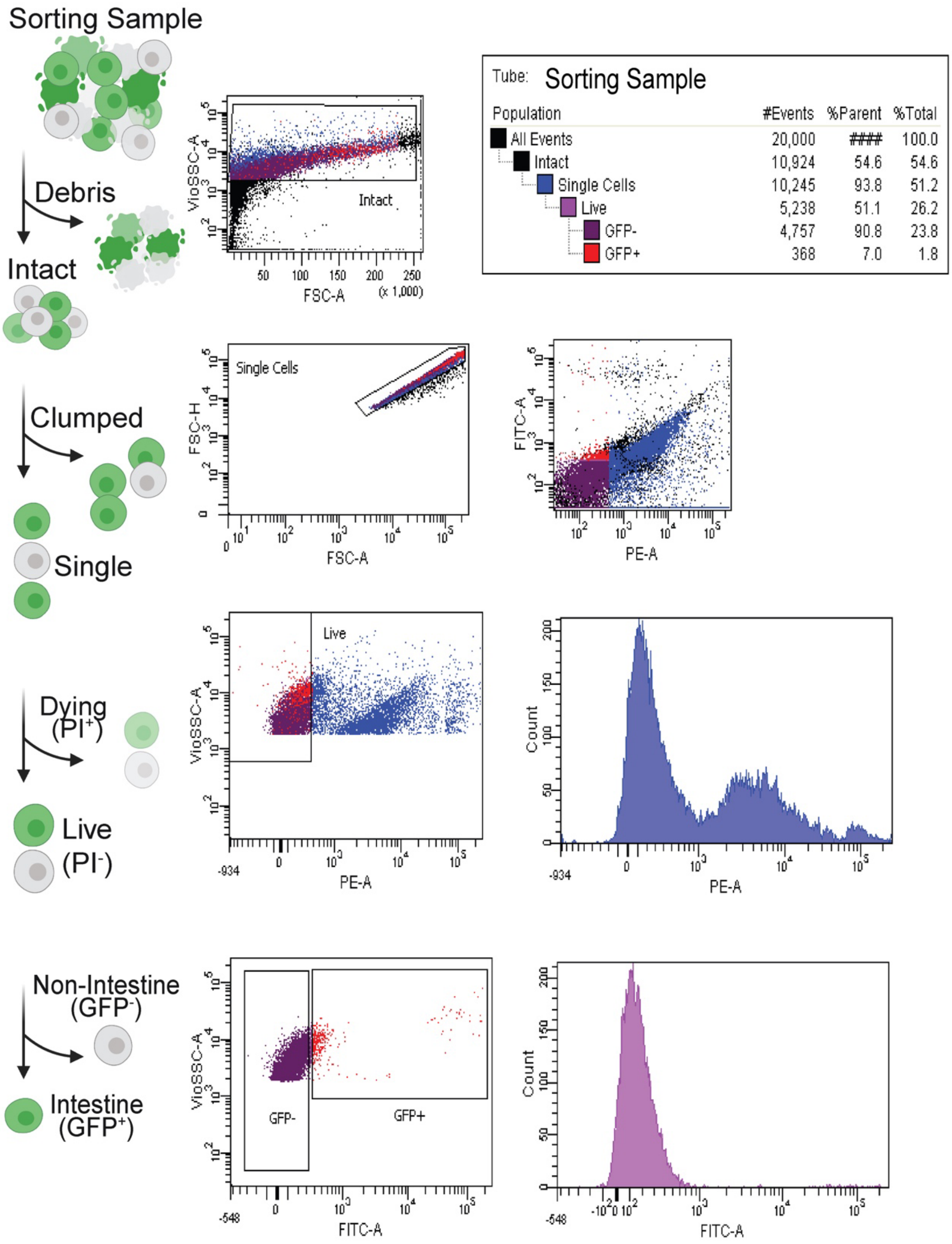


**Supplementary Figure 1. Gating strategy for Fluorescence Activated Cell Sorting (FACS) of embryonic *C. elegans* cells.** A mixed population of embryonic *C. elegans* embryos were dissociated into a single cell suspension using a combination of chemical and mechanical methods. Cells were then stained with Propidium Iodide (PI) for live/dead cell discrimination and run through a FACS Aria III cell sorter. Intestinal cells ($GFP^+$) and the all cells ($GFP^+$ and $GFP^-$) control group were acquired from an intact, single, and live cell population.

### Figure S2

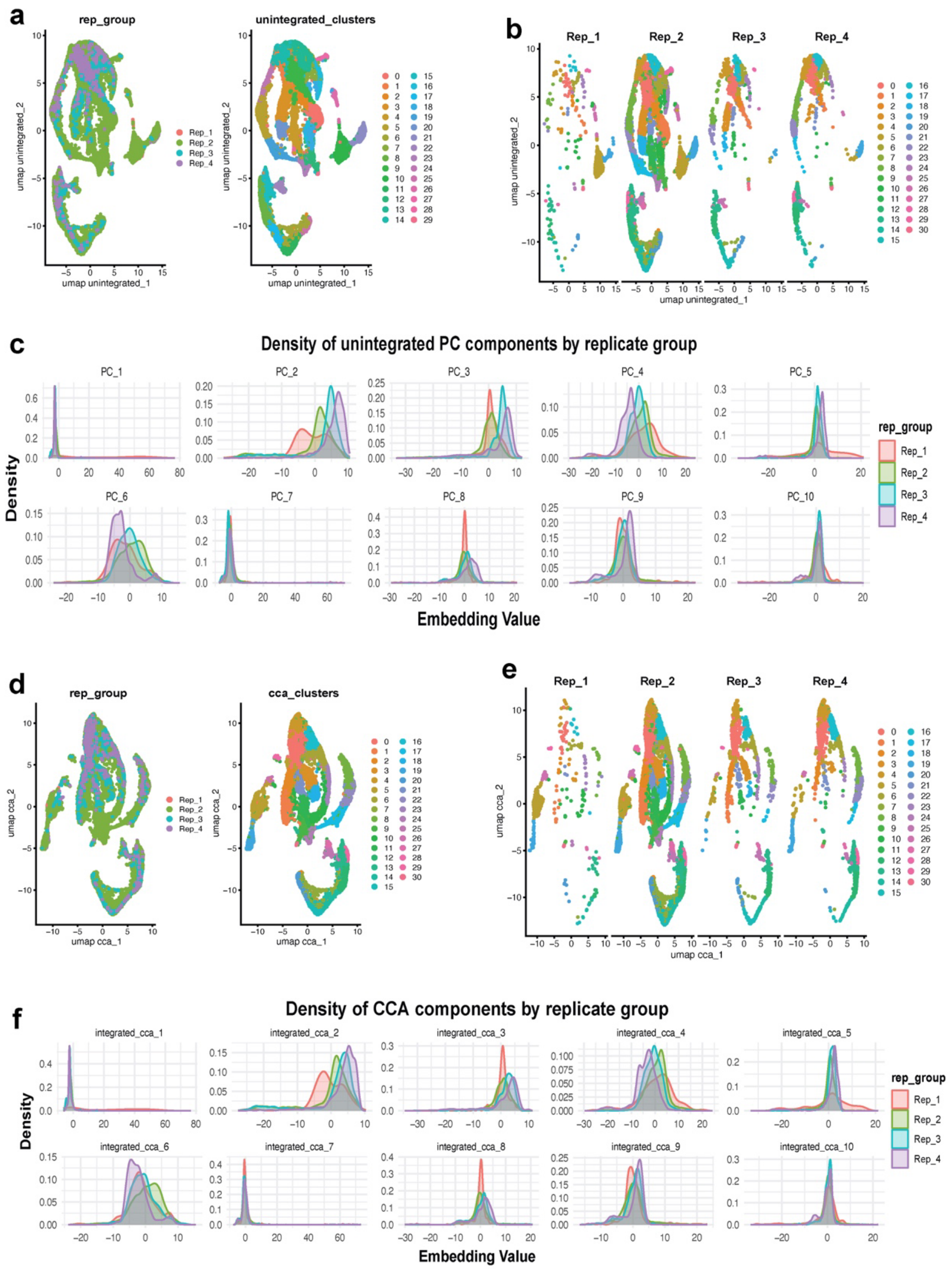


**Supplementary Figure 2. Diagnostic plots for embryonic *C. elegans* cells dataset pre and post integration of biological replicates.** Four biological replicates were collected for the all embryonic cells dataset. The data is broken up into pre-integration (**a-c**) and post-integration (**d-f**). **a)** UMAP embeddings of the scRNA-seq data from a mixed stage population of embryonic *C. elegans* cells with labeling of replicate (left) and identified clusters (right). **b)** UMAP embeddings of each of the four biological replicates split apart. **c)** Density histograms of the main principal component (PC) axis for visualization of data distribution by replicate, allowing visualization of batch effects. Figure panels **d–f** display the same types of information but post-integration of the dataset using the Anchor-based Canonical Correlation Analysis (CCA) method.

**Fig S3**

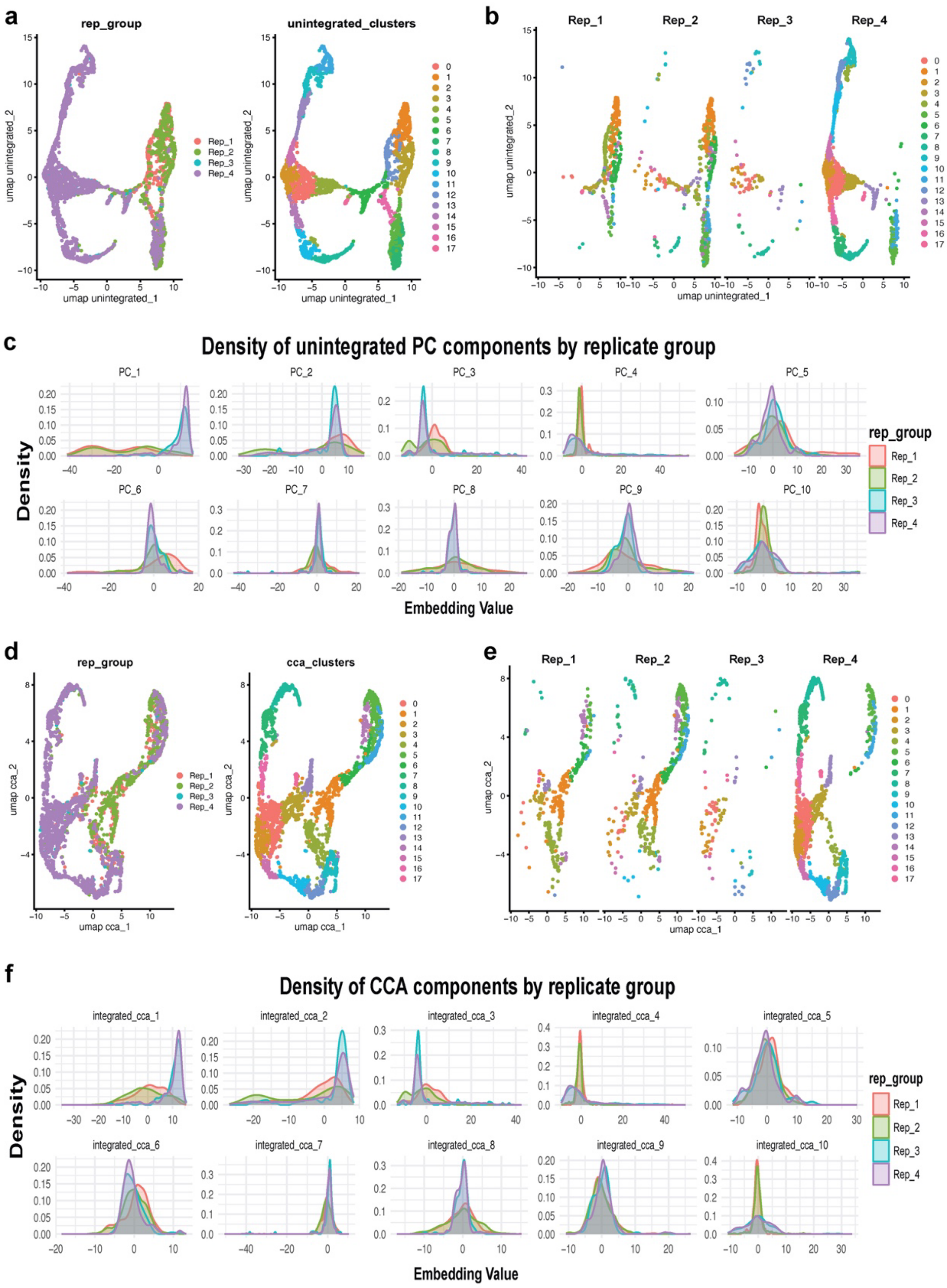


**Supplementary Figure 3. Diagnostic plots for embryonic *C. elegans* intestinal cells dataset pre- and post-integration of biological replicates.** Four biological replicates were collected for the sorted embryonic intestinal cells dataset. The data is broken up into pre-integration (**a-c**) and post-integration (**d-f**). **a)** UMAP embeddings of the scRNA-seq data from a mixed stage population of embryonic *C. elegans* intestinal cells with labeling of replicate (left) and identified clusters (right). **b)** UMAP embeddings of each of the four biological replicates split apart. **c)** Density histograms of the main principal component (PC) axis for visualization of data distribution by replicate, allowing visualization of batch effects. Figure panels **d–f** display the same types of information but post-integration of the dataset using the Anchor-based Canonical Correlation Analysis (CCA) method.

### Figure S4

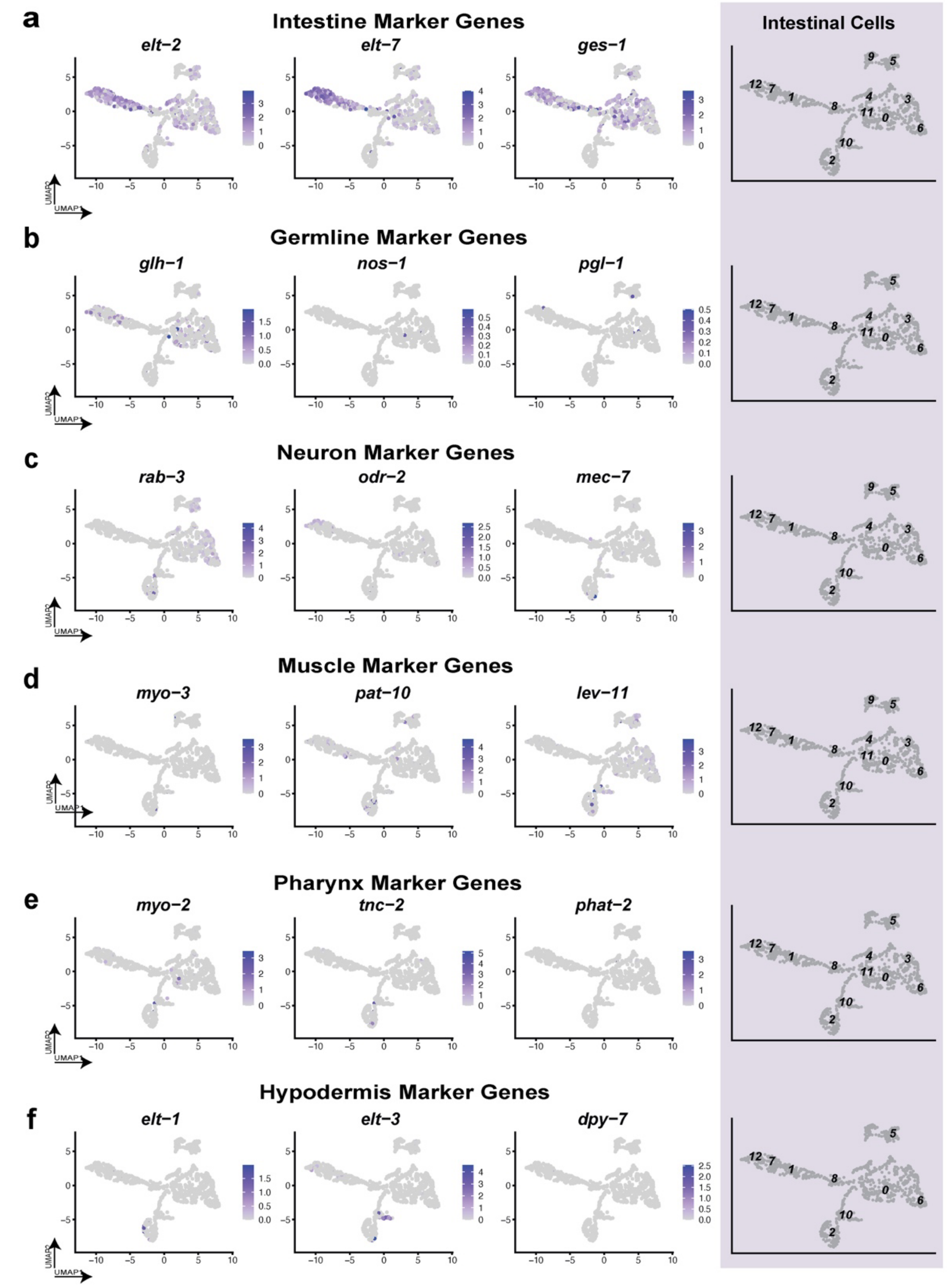


**Supplementary Figure 4. Tissue marker gene representation in sorted embryonic *C. elegans* intestinal cells.** (From left to right) Feature scatter plots displaying expression level and distribution within the UMAP embedding of tissue marker genes for the **a)** Intestine, **b)** Germline, **c)** Neurons, **d)** Muscle, **e)** Pharynx, and **f)** Hypodermis. The far right column in each row displays a UMAP embedding with the annotated cluster representation for each set of tissue marker genes.

### Figure S5

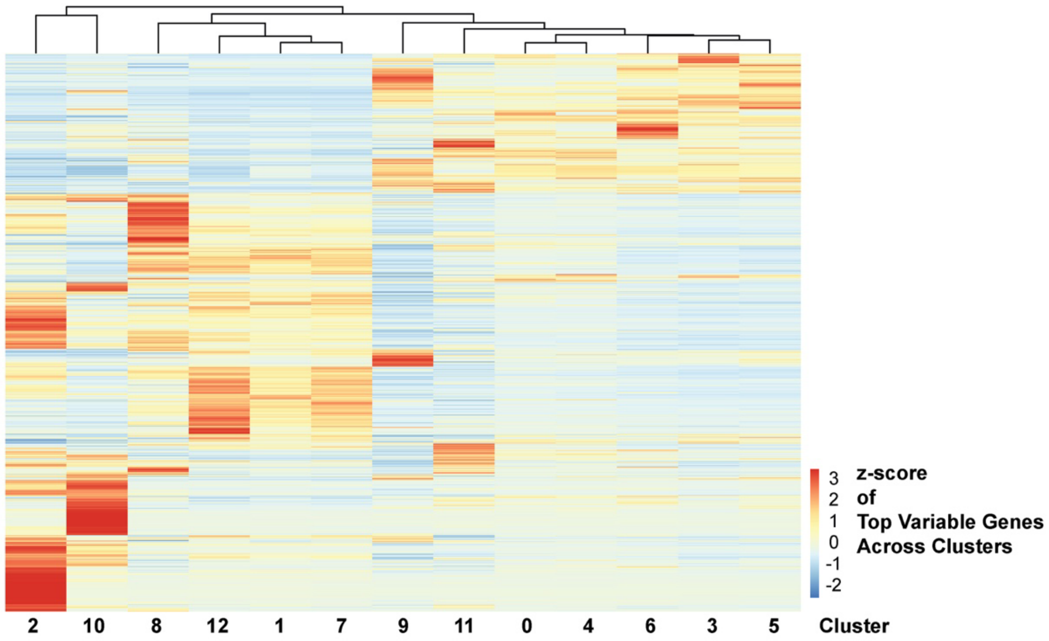


**Supplementary Figure 5. Top variable genes across identified *C. elegans* intestinal clusters.** A variance heatmap of gene expression, with identified clusters/cells as columns and genes as rows. Gene expression was aggregated at the cluster level, generating a matrix of average expression values (genes X clusters). For each gene, variance across clusters was calculated to quantify differences in mean expression between clusters. Genes with variance greater than 0.1 were retained and ranked by decreasing variance. This resulting subset is visualized as a heatmap after z-score scaling across clusters (row-wise normalization). Colors represent relative expression levels (scaled).

### Figure S6

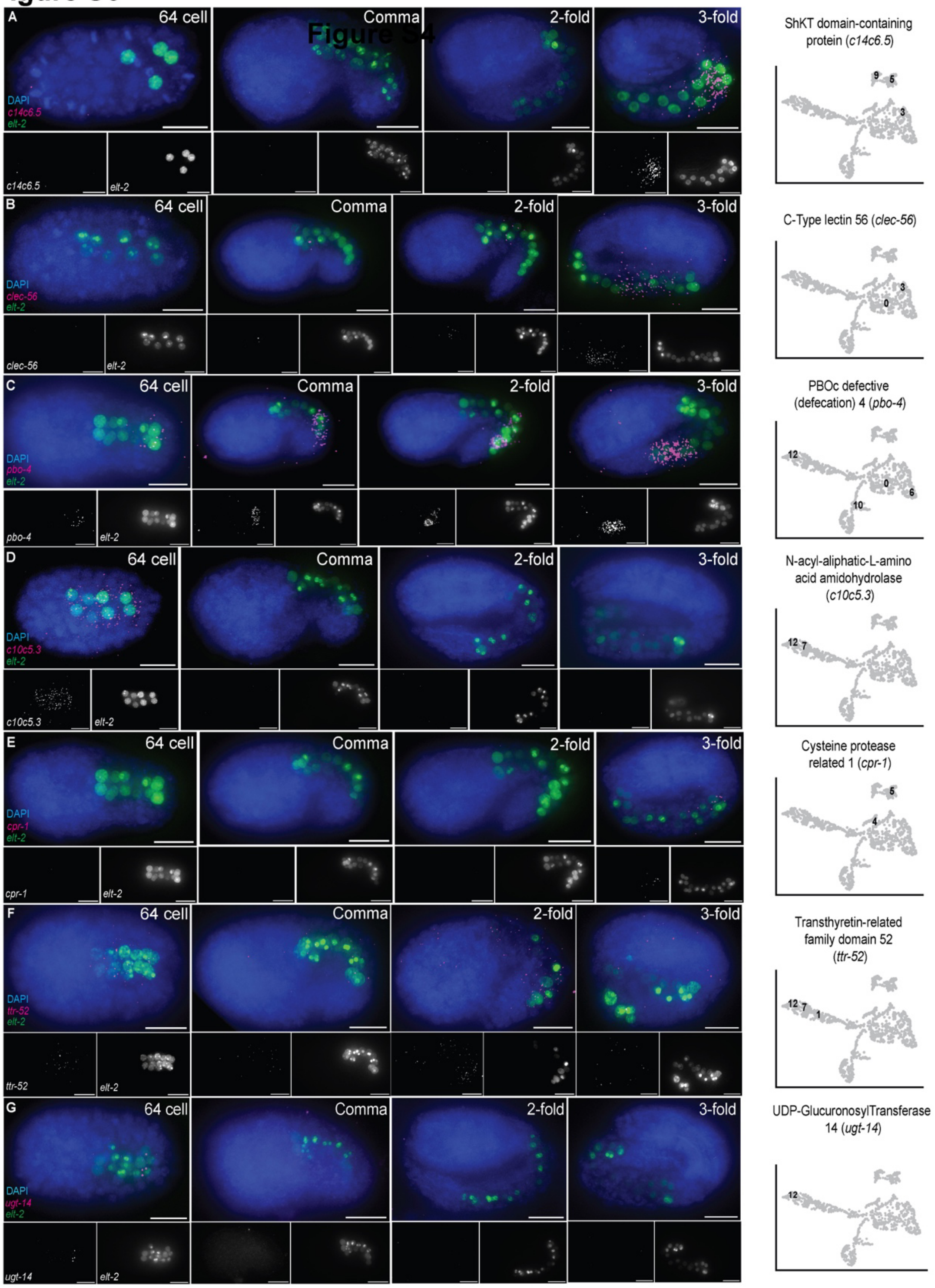


**Supplementary Figure 6. smiFISH targets evaluated for resolving *C. elegans* intestinal clusters (Part A).** The following transcripts can be found in Table 2 and were chosen for targeting with smiFISH to spatiotemporally validate intestinal cell clusters. From left to right are, representative microscopy images over developmental time for a given smiFISH target with corresponding feature plot showing expression across cells in the UMAP. Transcripts shown here include: *c14c6.5* (**A**), *clec-56* (**B**), *pbo-4* (**C**), *c10c5.3* (**D**), *cpr-1* (**E**), *ttr-52* (**F**), *ugt-14* (**G**).

### Figure S7

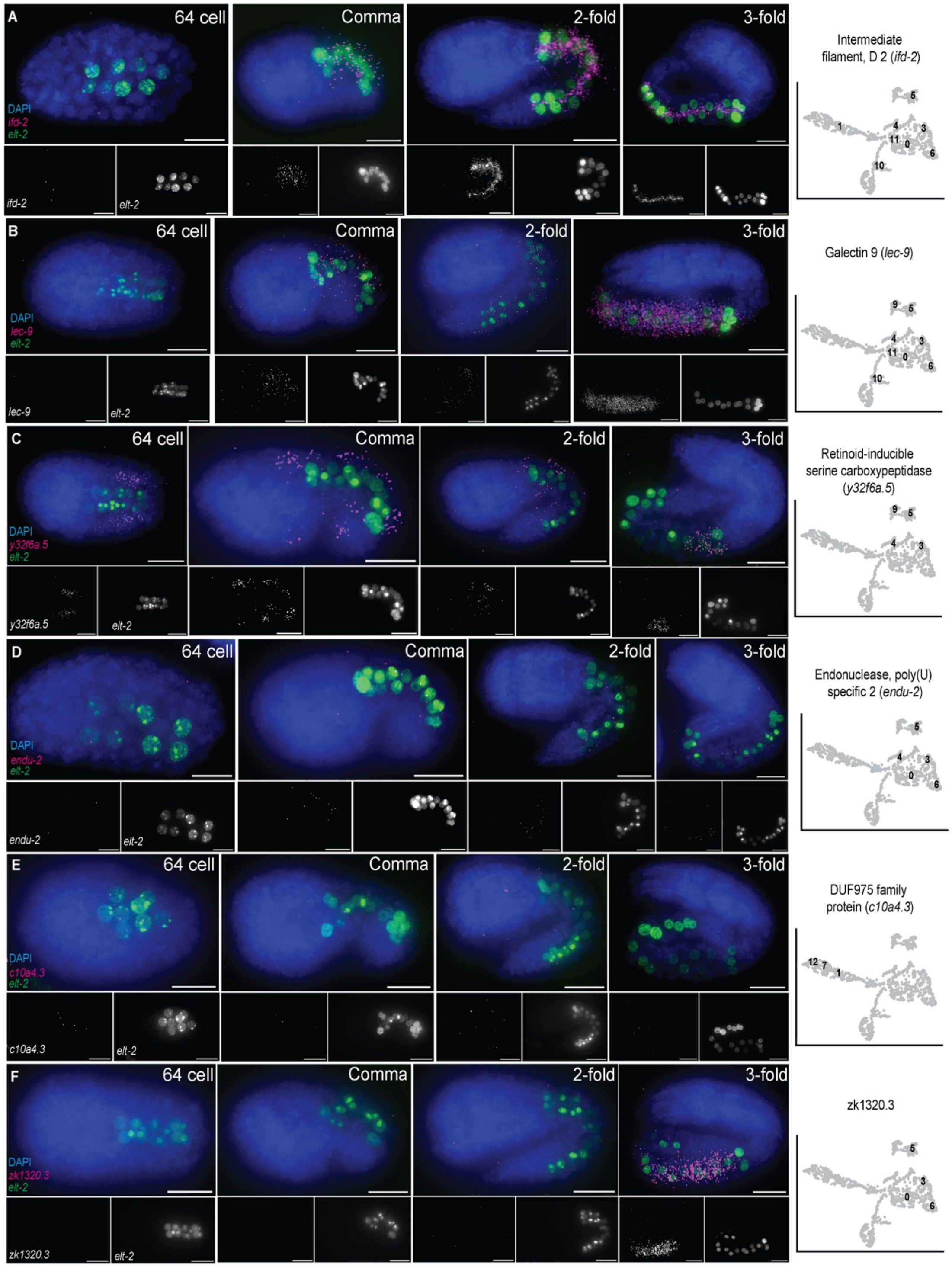


**Supplementary Figure 7. smiFISH targets evaluated for resolving *C. elegans* intestinal clusters (Part B).** The following transcripts can be found in Table 2 and were chosen for targeting with smiFISH to spatiotemporally validate intestinal cell clusters. From left to right are, representative microscopy images over developmental time for a given smiFISH target with corresponding feature plot showing expression across cells in the UMAP. Transcripts shown here include: *ifd-2* (**A**), *lec-9* (**B**), *y32f6a.5* (**C**), *endu-2* (**D**), *c10a4.3* (**E**), *zk1320.3* (**F**).

### Figure S8

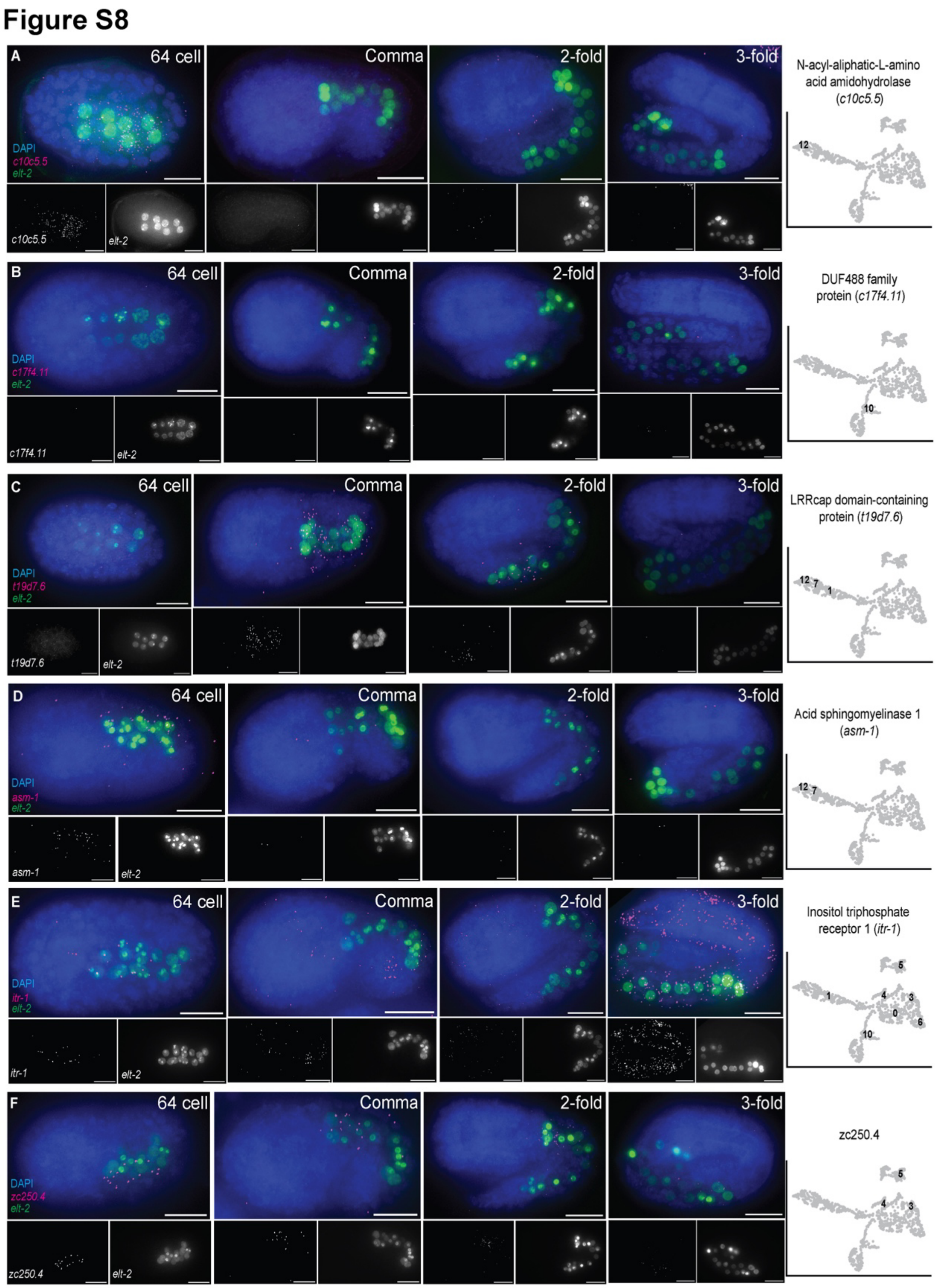


**Supplementary Figure 8. smiFISH targets evaluated for resolving *C. elegans* intestinal clusters (Part C).** The following transcripts can be found in Table 2 and were chosen for targeting with smiFISH to spatiotemporally validate intestinal cell clusters. From left to right are, representative microscopy images over developmental time for a given smiFISH target with corresponding feature plot showing expression across cells in the UMAP. Transcripts shown here include: *c10c5.5* (**A**), *c17f4.11* (**B**), *t19d7.6* (**C**), *asm-1* (**D**), *itr-1* (**E**), *zc250.4* (**F**).

Fig S9

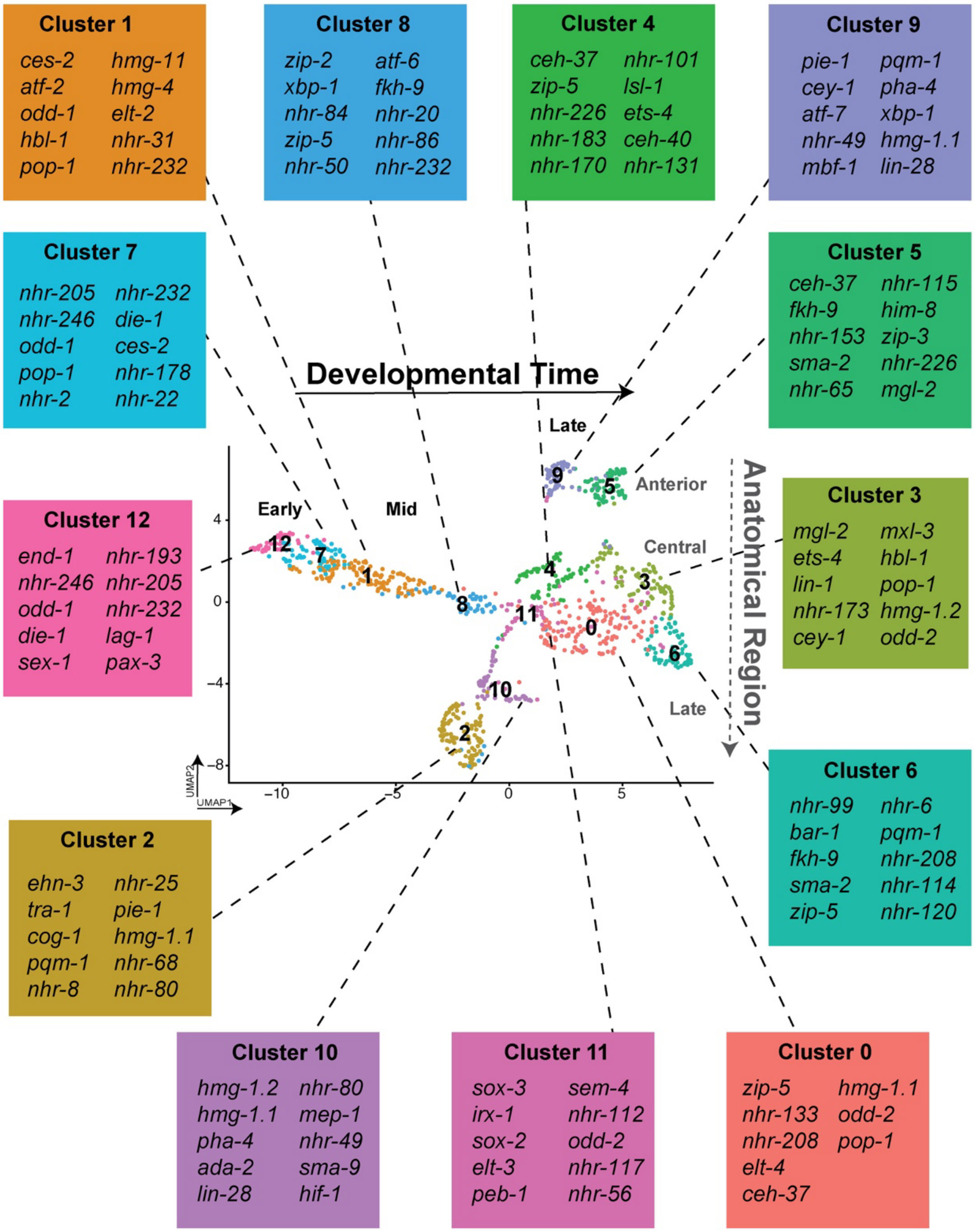


**Supplementary Figure 9. Top ten intestinal transcription factors per cluster.** UMAP projection of intestinal cell clusters annotated according to inferred developmental timing and anatomical organization within the intestine. For each intestinal cluster, the top 10 enriched transcription factors were pulled out from the cluster marker gene list by cross-referencing against the *C. elegans* transcription factor gene set referred to as wTF2.0 and known intestinal transcription factors pulled from the modEncode project.

**Fig S10**

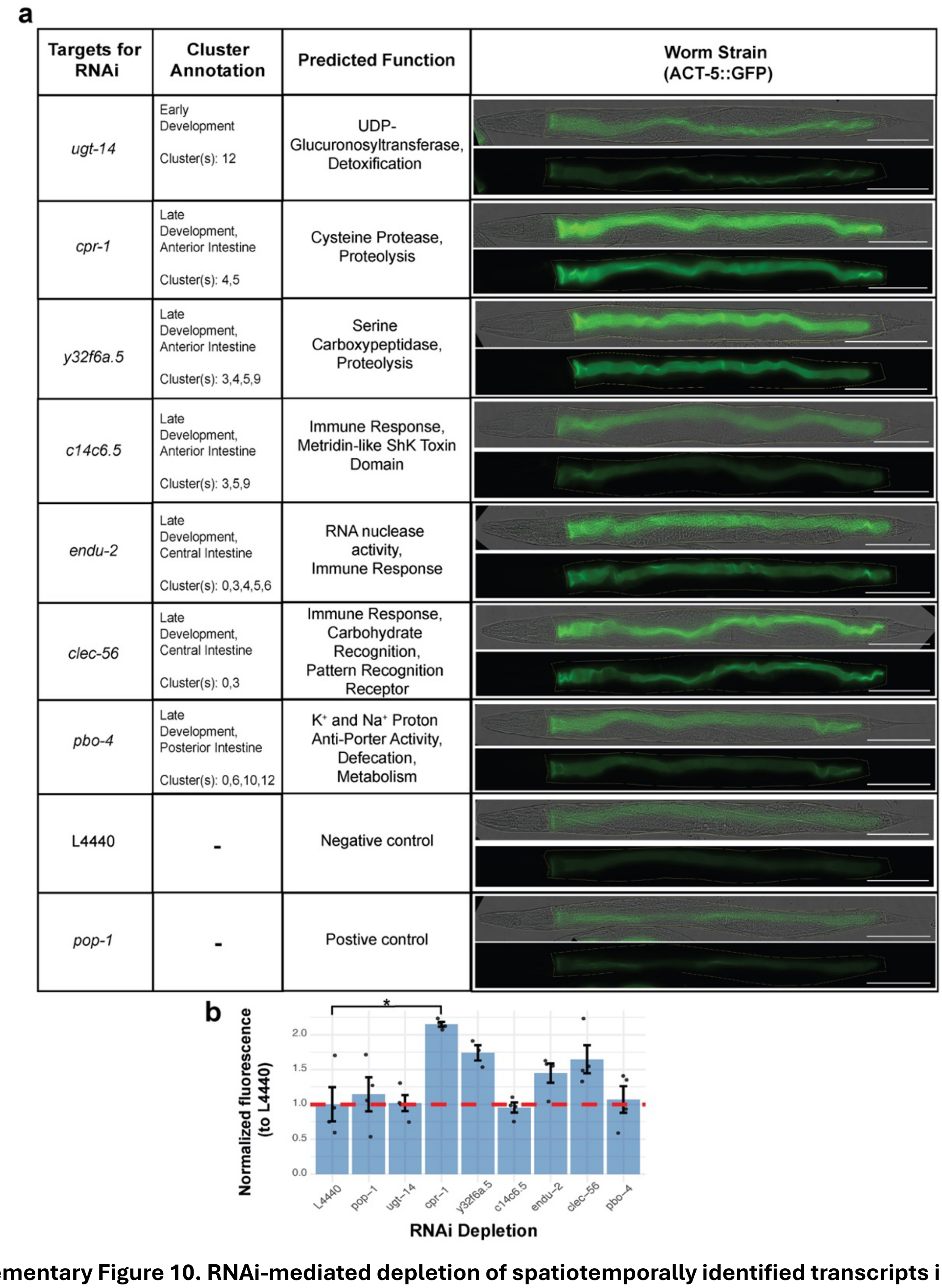


**Supplementary Figure 10. RNAi-mediated depletion of spatiotemporally identified transcripts in ERT60 worm strain.** (**a**) Representative brightfield and fluorescence images of L4 stage ERT60 (ACT-5::GFP) worms after RNAi depletion of spatiotemporally distinct candidate smiFISH targets (*ugt-14*, *cpr-1*, *y32f6a.5*, *c14c6.5*, *endu-2*, *clec-56*, and *pbo-4*), in addition to treatment with the both negative (L4440) and positive (*pop-1*) control RNAi strains. (**b**) Quantification of intestinal ACT-5::GFP fluorescence after RNAi treatment. Fluorescence intensity was measured in individual animals, background corrected and then normalized to the mean fluorescence of the negative control (L4440). Bars represent mean normalized fluorescence ± SEM, and black dots indicate individual animals. The red dashed line marks the normalized L4440 negative control (1.0). Statistical significance was determined by comparing each RNAi treatment to the negative control using two-sided Wilcoxon rank-sum test (*p-value<0.05).

**Fig S11**

**a**

| Targets for RNAi | Cluster Annotation | Predicted Function | Worm Strain (elt-2p::GFP; *cdc-25.1(rr31)*) |
|---|---|---|---|
| *ugt-14* | Early Development<br>Cluster(s): 12 | UDP-Glucuronosyltransferase, Detoxification | 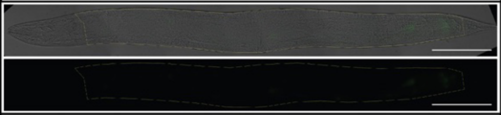 |
| *cpr-1* | Late Development, Anterior Intestine<br>Cluster(s): 4,5 | Cysteine Protease, Proteolysis | 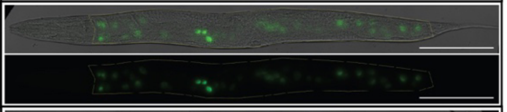 |
| *y32f6a.5* | Late Development, Anterior Intestine<br>Cluster(s): 3,4,5,9 | Serine Carboxypeptidase, Proteolysis | 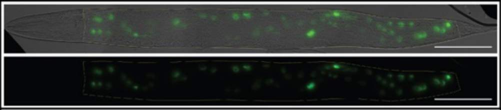 |
| *c14c6.5* | Late Development, Anterior Intestine<br>Cluster(s): 3,5,9 | Immune Response, Metridin-like ShK Toxin Domain | 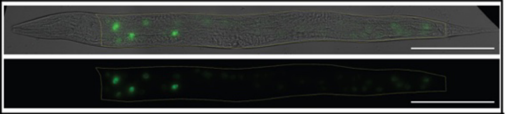 |
| *endu-2* | Late Development, Central Intestine<br>Cluster(s): 0,3,4,5,6 | RNA nuclease activity, Immune Response | 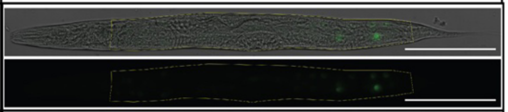 |
| *clec-56* | Late Development, Central Intestine<br>Cluster(s): 0,3 | Immune Response, Carbohydrate Recognition, Pattern Recognition Receptor | 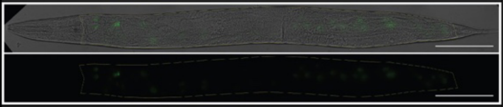 |
| *pbo-4* | Late Development, Posterior Intestine<br>Cluster(s): 0,6,10,12 | K⁺ and Na⁺ Proton Anti-Porter Activity, Defecation, Metabolism | 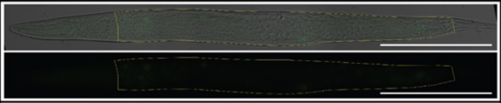 |
| L4440 | - | Negative control | 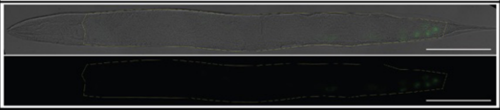 |
| *pop-1* | - | Postive control | 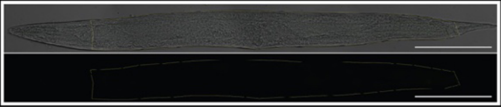 |

**b**

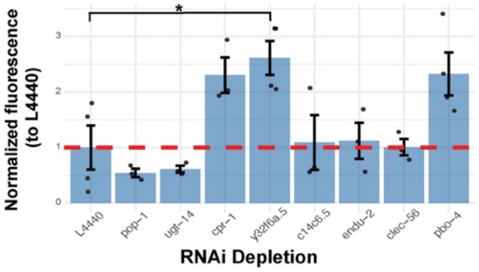


**Supplementary Figure 11. RNAi-mediated depletion of spatiotemporally identified transcripts in MR142 worm strain.** (**a**) Representative brightfield and fluorescence images of L4 stage MR142 (*elt-2p*::GFP; *cdc-25.1 (rr31)*) worms after RNAi depletion of spatiotemporally distinct candidate smiFISH targets (*ugt-14*, *cpr-1*, *y32f6a.5*, *c14c6.5*, *endu-2*, *clec-56*, and *pbo-4*), in addition to treatment with the both negative (L4440) and positive (*pop-1*) control RNAi strains. (**b**) Quantification of intestinal *elt-2p*::GFP fluorescence after RNAi treatment. Fluorescence intensity was measured in individual animals, background corrected and then normalized to the mean fluorescence of the negative control (L4440). Bars represent mean normalized fluorescence $\pm$ SEM, and black dots indicate individual animals. The red dashed line marks the normalized L4440 negative control (1.0). Statistical significance was determined by comparing each RNAi treatment to the negative control using two-sided Wilcoxon rank-sum test. One outlier from the *c14c6.5* RNAi treated group was excluded prior to analysis (*p-value<0.05).
